# Deep Learning-Guided Holotomography Reveals Early Structural Remodelling During Pluripotency Exit

**DOI:** 10.64898/2026.04.23.720508

**Authors:** Hoewon Park, Geon Kim, Jeongwon Shin, Seo-Hyun Kim, Eui-been Hwang, Minji Kim, Taewoong Hwang, Arim Lim, Gyungtae Yoon, Jisu Park, Young-woo Jeon, Nam-Shik Kim, YongKeun Park, Ki-Jun Yoon

## Abstract

Real-time assessment of human pluripotent stem cell (hPSC) quality is critical for reproducibility and safety in regenerative medicine, yet current methods are invasive, labor-intensive, or highly operator-dependent. We present DeepHOPE (Deep-learning-guided Holotomography for Pluripotency Evaluation), a non-invasive, automatizable, and statistics-driven platform that integrates three-dimensional (3D) refractive index imaging with deep learning to assess pluripotency. DeepHOPE performs robustly across diverse contexts, including germ-layer differentiation, retinoic acid–induced differentiation, and mid-reprogramming cultures, enabling streamlined cell production workflows and improving the efficiency of midbrain dopaminergic neuron differentiation through informed colony selection. Mechanistically, DeepHOPE detects minute colony-scale topological changes that precede molecular loss of pluripotency. These early changes are associated with rapid F-actin remodeling, including apical-to-basal redistribution during early differentiation. Consistent with a functional role for cytoskeletal regulation in state transitions, sustained reduction of actomyosin tension decreases pluripotency, identifying cytoskeletal dynamics as an upstream determinant of early pluripotency exit.

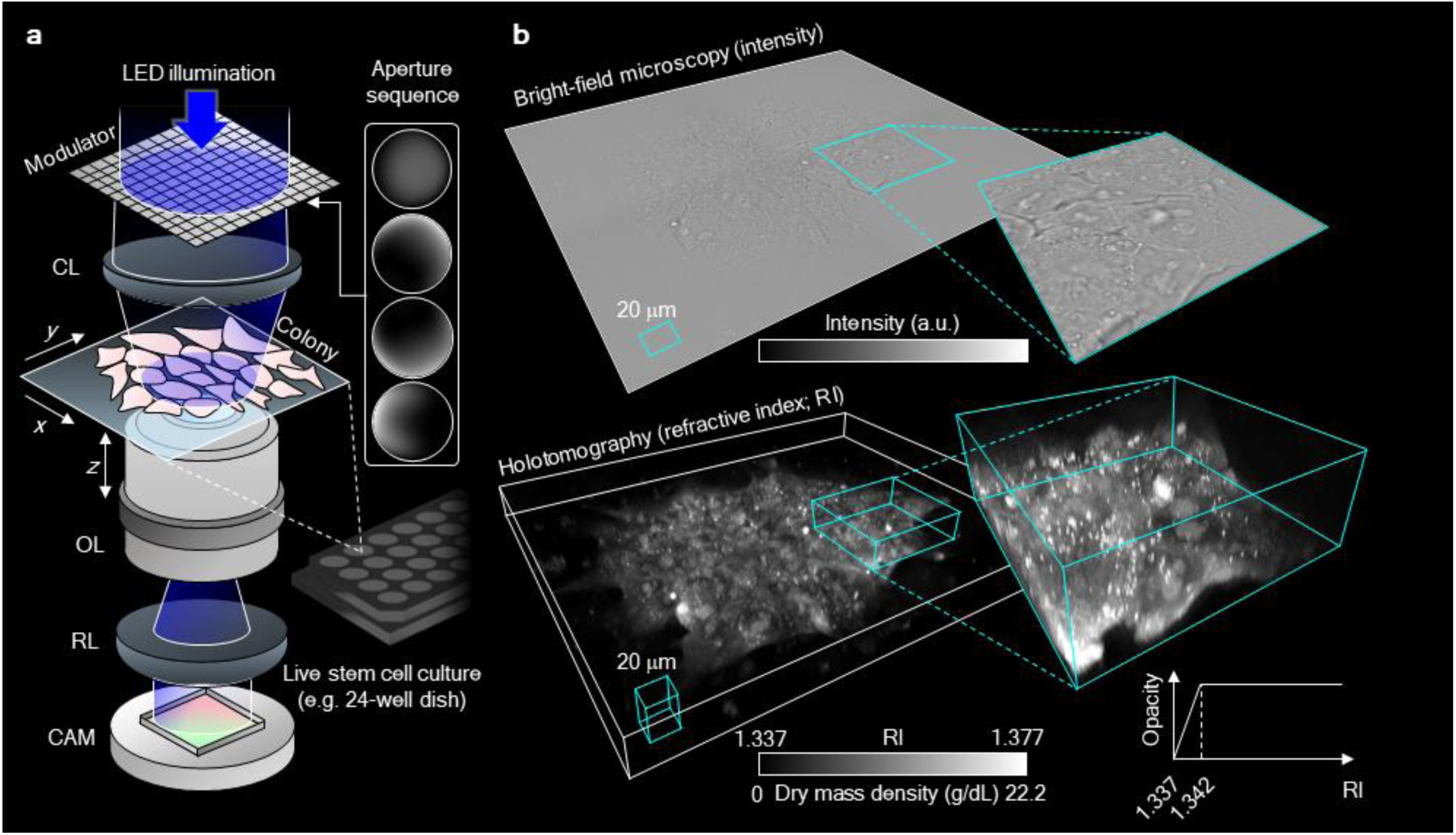

## Introduction

Human pluripotent stem cells (hPSCs), comprising human embryonic stem cells (hESCs) and human induced pluripotent stem cells (hiPSCs), have revolutionised the ability to model functional human tissues in a patient-specific manner, directed by specific signaling molecules and growth factors^1–3^. More recently, the development of hPSC-derived organoids—capable of recapitulating the three-dimensional (3D) heterogeneity of native tissues— has intensified the demand for robust, high-quality hPSC sources across applications in disease modelling, drug discovery, and precision medicine^4,5^. Despite this progress, the reliable production and maintenance of hPSCs remain technically challenging, particularly in sustaining the undifferentiated state and ensuring reproducible differentiation outcomes^6,7^. Pluripotency is highly sensitive to experimental variables, including culture conditions, passaging methods, and user-dependent handling protocols^8,9^. In hiPSCs, inter- and intra-batch variability further arises from donor cell status, reprogramming method, and colony selection criteria^10,11^. These challenges highlight the urgent need for standardised, quantitative quality-control frameworks.

Conventional approaches to assessing pluripotency typically rely on molecular assays that quantify RNA or protein expression of core pluripotency markers such as PluriTest^12,13^, yet these methods are inherently labor-intensive, destructive, and incompatible with continuous monitoring of live cells. Fluorescence imaging–based hPSC profiling has also been employed but is limited by the requirement for fixation or significant phototoxicity during *in situ* assessment of pluripotency^14,15^. Thus, conventional label-free imaging^16–19^ has been explored as non-invasive surrogates for cellular identity, leveraging structural features as proxies for pluripotency. These methods, often augmented by computational image analysis, have shown potential in distinguishing undifferentiated from differentiating colonies based on features such as nucleolar prominence, cytoplasmic sparsity, and colony boundary integrity^16,17,20,21^. However, their interpretive power remains limited; morphological classifiers exhibit reduced sensitivity in borderline cases, and the intrinsically qualitative, two-dimensional nature of these modalities hampers consistent, high-resolution evaluation^22–25^.

To address these limitations, we adopted holotomography (HT)^26–31^, a three-dimensional quantitative phase imaging technique that reconstructs the refractive index (RI) distribution of biological samples by numerically solving the inverse light-scattering problem. This method yields sub-micrometer resolution (∼150 nm) while operating under low photon doses, thereby minimizing phototoxicity and enabling rapid acquisition; its technical characteristics and biomedical applications have been reviewed in detail^29^. RI tomograms provide rich, quantitative information on intracellular architecture, offering substantially greater spatial resolution and contrast than traditional label-free microscopy, and making HT ideally suited for longitudinal analysis of live hPSC colonies. Crucially, the integration of HT with machine learning further expands its analytical utility. By coupling high-resolution RI data with deep learning algorithms, complex morphological features can be quantitatively linked to biological states that would otherwise require destructive molecular assays^28–30^. This approach enables non-invasive, real-time inference of cellular identity, offering a powerful and scalable alternative for stem cell quality control.

Here, we present DeepHOPE (Deep-learning-guided HOlotomography for Pluripotency Evaluation), a robust framework that integrates 3D HT imaging with deep neural network (DNN)-based classification to assess pluripotency at high spatial resolution (Extended Data Fig. 1). DeepHOPE is inspired by previous works that have computationally related light scattering or HT-trackable morphologies to the phenotypic status including stem cell physiology^32–35^. DeepHOPE performs spatially resolved classification across hPSC colonies by identifying subtle morphological signatures indicative of early differentiation into the three germ layers. We trained and validated the DNN using curated datasets from standard differentiation protocols and demonstrated its applicability across diverse experimental conditions.

In addition, DeepHOPE-based interpretability and quantitative 3D refractive index feature analysis identified colony flattening as an early and robust structural signature of pluripotency exit, detectable before overt molecular loss of pluripotency markers. This transition is accompanied by rapid F-actin remodeling and early mechanical changes, indicating that cytoskeletal dynamics are a key upstream driver of early colony-state transitions. Our results underscore the potential of DeepHOPE as a non-invasive, real-time solution for quantitative pluripotency assessment and hiPSC quality control, enabling scalable and standardised stem cell manufacturing workflows while providing mechanistic insight into the early stages of pluripotency exit.

## Results

### Quantitative 3D characterisation of hPSC colonies using holotomography

Prior to classification, HT itself offers substantial advantages for label-free structural interrogation of hPSC colonies over time. Unlike bright-field microscopy, HT provides superior spatial resolution and quantitative contrast based on intrinsic RI variations, allowing visualisation of both cellular and subcellular architecture without exogenous labels (Fig. 1a, b)^32^. The resulting RI tomograms revealed consistent structural features across the colony, including sharp boundaries, dense cellular cores, and layered morphology, which recapitulate known 3D epithelial-like arrangements seen in fluorescence-based studies (Fig. 1c,d)^14,33^. In large colonies exceeding 500 µm in diameter, we observed distinct spatial patterns: central cells appeared densely packed with relatively low RI, while peripheral cells were radially aligned and exhibited higher RI values (Fig. 1c and Extended Data Fig. 2). Notably, without nuclear or cytoskeletal staining, HT enabled identification of nucleoli, nuclear envelopes, intercellular gaps, lipid droplets, and lamellipodia-like membrane protrusions displaying dynamic ruffling behaviour (Fig. 1d and Extended Movie 1). Mitotic chromosomes were also readily discernible indicating the cell cycle stage in live colonies (Fig. 1e and Extended Movie 2). These findings highlight the sensitivity of HT for capturing fine-scale morphodynamic features in live hPSCs, laying the foundation for image-based, label-free assessment of pluripotency and differentiation potential.

**Figure. 1.**
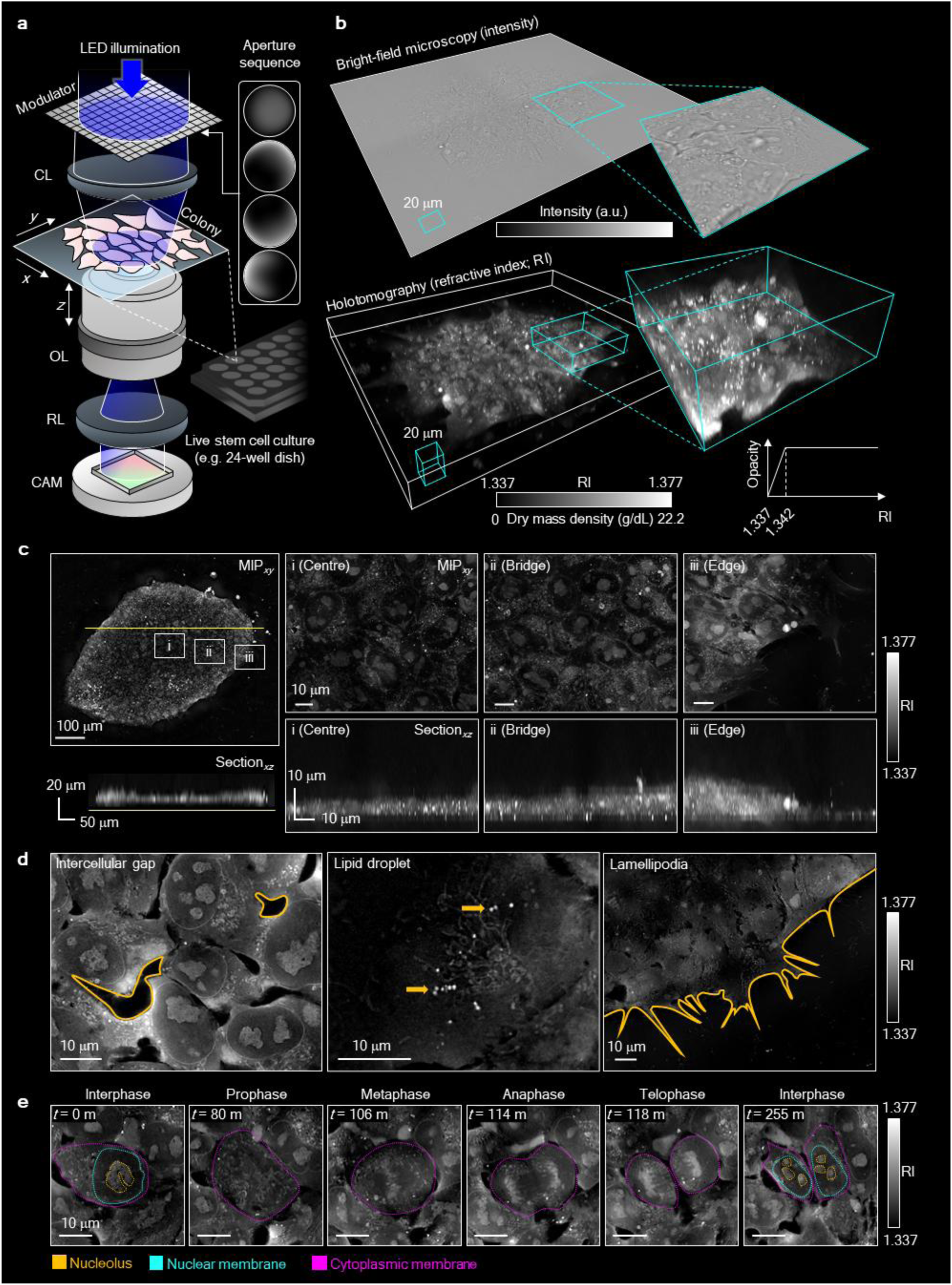
High-resolution HT imaging of live hPSC colonies reveals subcellular and topographical features. (**a**) Schematic of the low-coherence HT system. The setup uses modulated LED illumination and sequential aperture patterns to acquire light scattering profiles across multiple focal planes (*z*-axis), enabling reconstruction of 3D RI tomograms. CL, condenser lens; OL, objective lens; RL, relay lens; CAM, camera. (**b**), Comparison of bright-field microscopy (top) and HT imaging (bottom) of an undifferentiated GM25256 hPSC colony. HT provides enhanced spatial contrast and RI-based quantification of intracellular composition. Insets show higher magnification views of selected regions. (**c**) HT imaging of a large hPSC colony. Maximum intensity projection (MIP) and vertical cross-sectional views reveal spatial heterogeneity in colony architecture, including a densely packed centre (i), an elevated bridging region (ii), and tilted cells at the colony periphery (iii). (**d**) Representative horizontal sections highlighting key subcellular structures observed without labelling: intercellular gaps (left), lipid droplets (middle; yellow arrows), and lamellipodia-like peripheral protrusions (right). (**e**) Time-lapse HT imaging capturing distinct phases of the mitotic cycle in live hPSCs. Subcellular components, including nucleoli (yellow), nuclear membranes (cyan), and cytoplasmic membranes (magenta), are visualised using label-free RI segmentation.

### Development and validation of DeepHOPE for pluripotency assessment

We next constructed the deep neural network (DNN) to complete the DeepHOPE framework. Through this DNN, we sought to capture the structural signatures of early-stage differentiation of hPSCs. Spontaneous differentiation, commonly triggered by suboptimal culture conditions or mechanical stress during passaging, often manifests as subtle and heterogeneous changes in colony structure. We hypothesised that the morphological transitions accompanying early lineage commitment could mirror those in spontaneous differentiation and serve as reliable, image-based indicators of pluripotency loss. To construct a training dataset, we induced directed differentiation of a standard hiPSC line (GM25256) towards the three germ layers—endoderm, mesoderm, and ectoderm—and acquired time-course HT images at 6, 12, 24, and 48 hours post-induction (Fig. 2a-c and Extended Data Fig. 3a-g). GM25256 colonies under ectoderm-directed differentiation were measured at an additional time point of 72 hours post-induction, to reflect the longer duration of ectoderm commitment. These data, representing progressive morphological shifts away from the pluripotent state^34–36^, were used to define the “differentiating” class. “Undifferentiated” colonies maintained under control conditions were used as the reference class.

**Figure. 2.**
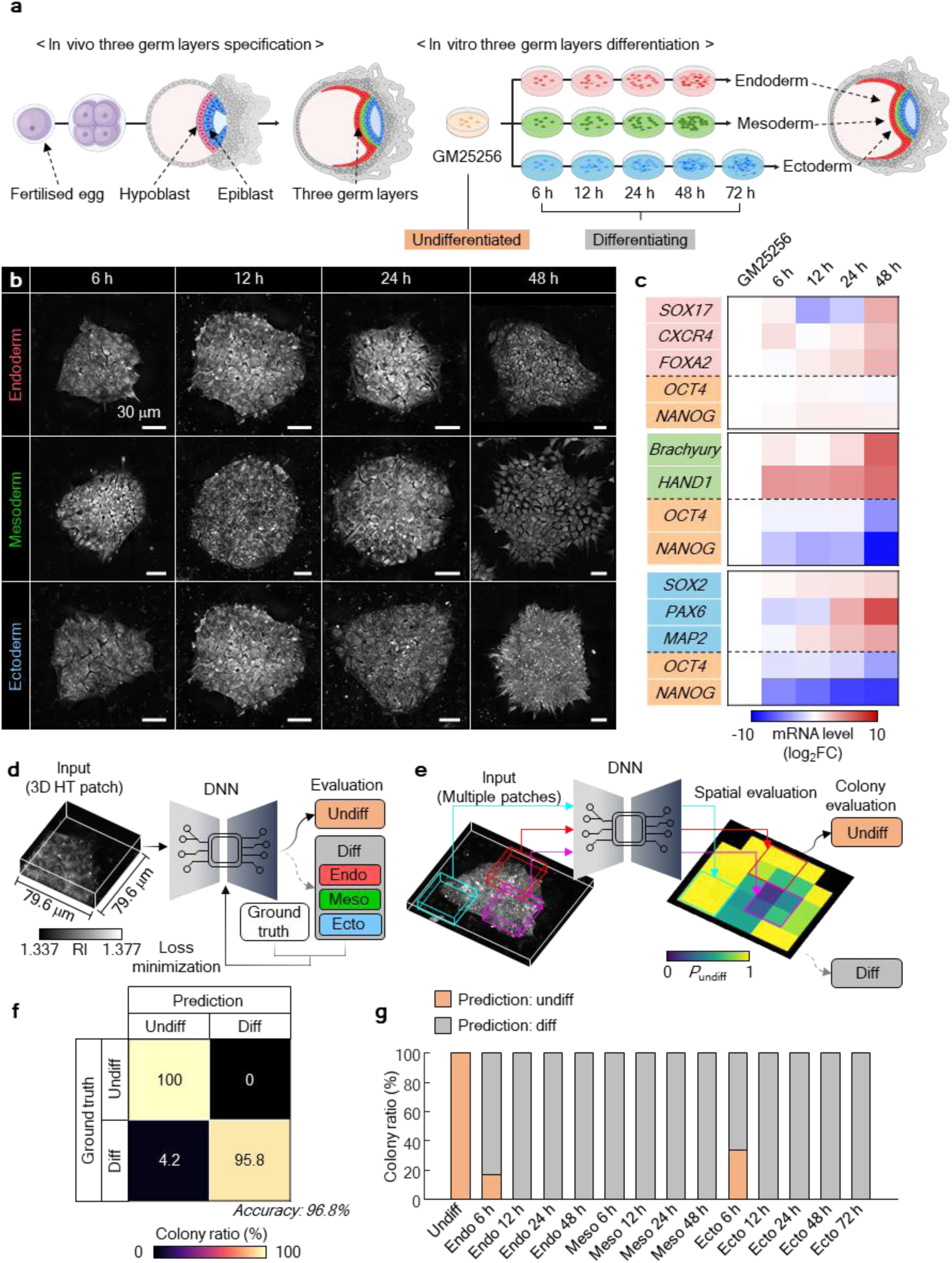
Development and validation of DeepHOPE using directed differentiation of hPSC colonies. **(a)** Trilineage differentiation experiment for generating training data for DeepHOPE. **(b)** Representative HT images of hPSC colonies collected at 6, 12, 24, and 48 hours following germ layer-specific induction. Images are shown as maximum intensity projections along the *z*-axis. **(c)** Transcriptomic profiling of corresponding colonies reveals upregulation of lineage-specific markers (red, endoderm; green, mesoderm; blue, ectoderm) and concurrent downregulation of pluripotency genes (orange: OCT4, NANOG). **(d)** Schematic of the patch-wise DeepHOPE training workflow. **(e)** Schematic of the colony-level DeepHOPE inference process. Prediction probabilities across patches are aggregated to derive colony-level pluripotency status. **(f)** Confusion matrix showing colony-level classification accuracy in a blind test, achieving 96.8%. **(g)** Rates of colony-level DeepHOPE prediction across time points and differentiation lineages. *n* = 90 colonies for total groups.

To accommodate colony-to-colony variability in size, shape, and differentiation patterns, we designed a spatially localised classification framework (see Extended Data Fig. 4 for design). Each HT image was divided into fixed-size patches (79.6 μm × 79.6 μm), and a DNN was trained to classify each patch as either “undifferentiated” or “differentiating” (Fig. 2d and Supplementary Table 1). During inference, DeepHOPE integrates patch-level DNN outputs to produce a colony-level evaluation (Fig. 2e). To be specific, each patch-level output is the probability of classification into an “undifferentiated” patch denoted *P*_undiff_. Patch-wise *P*_undiff_ values are assembled to form a spatial *P*_undiff_ map and averaged to yield the colony-level *P*_undiff_. Each colony is evaluated “undifferentiated” if colony-level *P*_undiff_ is higher than 0.5 and “differentiating” otherwise. DeepHOPE achieved 96.8% classification accuracy on a blind test dataset (Fig. 2f). Notably, misclassifications were primarily observed at the earliest (6 hours) differentiation time point, when morphological differences remained minimal (Fig. 2g).

### Molecular validation of DeepHOPE across diverse human PSC status and backgrounds

To evaluate the broader applicability of DeepHOPE for detecting pluripotency loss, we first tested its performance in a chemically induced differentiation model using all-trans retinoic acid (RA), a well-established inducer of neural and pancreatic lineage specification in hPSCs^37–39^. In GM25256 hiPSC colonies treated with RA, mRNA levels of OCT4 and NANOG declined significantly after 48 hours, whereas earlier time points (12 and 24 hours) showed only minor reductions (Extended Data Fig. 5a). To capture corresponding morphological shifts, we performed time-course HT imaging of live colonies exposed to RA for up to 96 hours (Fig. 3a and Extended Data Fig. 5b-e). To relate DeepHOPE evaluation with molecular pluripotency, colonies were fixed and subjected to immunocytochemistry (ICC) immediately after HT acquisition for key pluripotency markers: OCT4, SSEA4, and NANOG (Fig. 3b). Colony-level *P*_undiff_ showed strong concordance with marker expression, yielding clear stratification of fluorescence intensities when colonies were binned according to *P*_undiff_ ranges (Fig. 3c). The rate of colonies evaluated “undifferentiated” by DeepHOPE was steeply reduced to 6.25% at 12 hours after RA treatment and sustained until 96 hours (Fig. 3d). To assess generalizability across genetic backgrounds, we further applied the same RA protocol to two additional hPSC lines (H9 and KOLF2.1J). Without RA treatment, 90.0% (H9) and 92.6% (KOLF2.1J) of colonies were classified as undifferentiated. After 12 hours of RA exposure, these proportions dropped to 49.7% and 57.4%, respectively, and further declined to 18.9% and 11.4% after 24 hours (Extended Data Fig. 5f-i). These results demonstrate that DeepHOPE robustly generalises across hPSC lines and sensitively detects RA-induced pluripotency loss in a time-dependent manner, as well as pluripotency loss beyond the conditions covered by the training data.

**Figure. 3.**
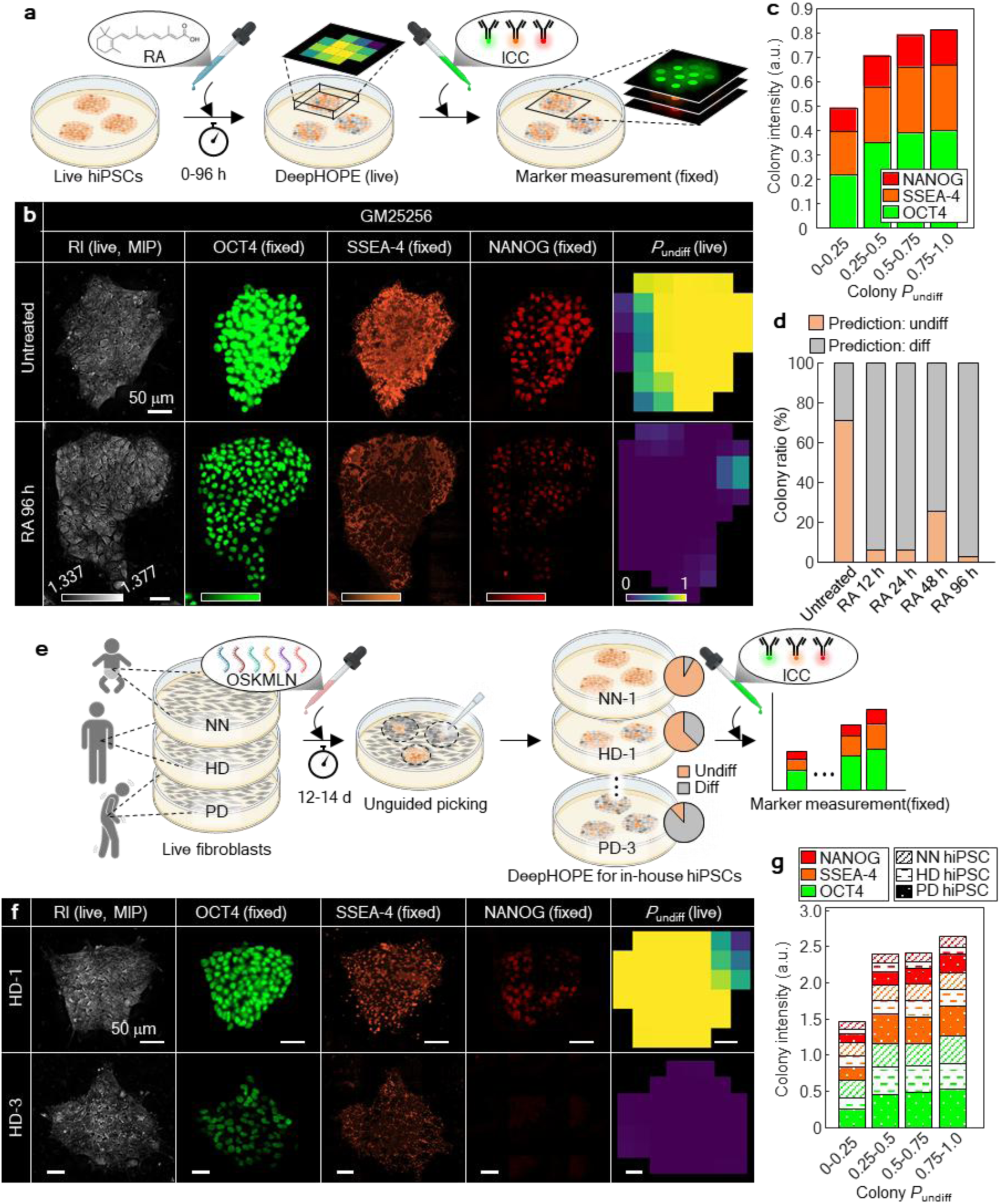
Validation and generalisation of DeepHOPE across RA-induced differentiation and reprogrammed hiPSCs. **(a)** Schematic of retinoic acid (RA)-induced spontaneous differentiation. Live-cell HT measurement was followed by fixation and staining with pluripotency marker **(b)** Representative HT (live) and fluorescence (fixed) images showing pluripotency marker expression (OCT4, SSEA4, NANOG) at baseline and following 48 h or 96 h RA exposure. The rightmost panels show DeepHOPE-derived spatial maps of *P*_undiff_ based on live HT images. Scale bar = 50 μm. **(c)** Stacked bar plot of marker fluorescence intensities across ranges of colony *P*_undiff_ from RA-treated hPSC colonies. **(d)** Rates of colony-level DeepHOPE prediction across time-courses after RA treatment. **(e)** Schematic of demonstrating DeepHOPE on fibroblasts-derived hiPSCs from three donors—a neonate (NN), healthy adult donor (HD), and Parkinson’s disease patient (PD). **(f)** Representative HT (live) and fluorescence (fixed) images from hiPSC colonies with high (top) and low (bottom) marker expression. The rightmost panels show DeepHOPE-derived *P*_undiff_ maps based on the live HT images. Scale bar = 50 μm. **(g)** Stacked bar plot of marker fluorescence intensities across ranges of colony *P*_undiff_ from reprogrammed hiPSC colonies.

We next evaluated whether DeepHOPE could assess pluripotency states of in-house hiPSCs isolated by unguided selection after fibroblast reprogramming. Reprogramming efficiency is known to vary substantially depending on donor cell status, delivery of reprogramming factors, and culture conditions, often resulting in a heterogeneous mixture of fully and partially reprogrammed colonies^40–42^. Thus, colony-level evaluation of live pluripotency is particularly valuable for personalised hiPSC applications. To test this, we analysed a panel of ten in-house hiPSC lines reprogrammed from fibroblasts derived from three donors of distinct biological backgrounds: a healthy adult (HD), a Parkinson’s disease patient (PD), and a neonate (NN). The hiPSC lines (3 from HD, 3 from PD, and 4 from NN) were randomly selected without prior screening for pluripotency. Each colony was imaged live using HT, followed by fixation and ICC to measure expression of OCT4, SSEA4, and NANOG (Fig. 3e,f). Across all lines, DeepHOPE-derived *P*_undiff_ showed a positive correlation with fluorescence intensities, supporting the framework’s applicability across diverse hiPSC backgrounds (Fig. 3g and Extended Data Fig. 6b). By contrast, an alternative workflow that replaces HT imaging with BF imaging showed significantly lower performance in evaluating reprogrammed hiPSCs (Extended Data Fig. 6a,c).

### DeepHOPE enables real-time selection of functionally competent hiPSC Colonies for personalised stem cell therapy

Because the DeepHOPE assessment can be repeatedly applied to live hPSC colonies (Extended Movie 3), it can serve a unique role in real-time monitoring of hiPSC reprogramming. We experimentally validated this hypothesised advantage in mid-reprogramming cultures. Specifically, neonatal fibroblasts were subjected to mRNA-based reprogramming for 12–14 days, after which regions containing visibly crowded cells were examined with DeepHOPE analysis to identify nascent hiPSC colonies (Fig. 4a). Fixation and ICC for pluripotency markers—OCT4, SSEA4, and NANOG—were performed immediately after live imaging to determine the molecular pluripotency states of the cells (Fig. 4b). The spatial distribution of DeepHOPE-derived *P*_undiff_ showed strong concordance with marker fluorescence intensities (Fig. 4c), supporting the utility of DeepHOPE for localizing high-quality hiPSC colonies. Quantitative analysis of local fluorescence intensity further confirmed a positive patch-level correlation with DeepHOPE-derived *P*_undiff_, although some variance was observed—likely reflecting biological asynchrony, fixation artifacts, or immunolabeling inefficiency. This highlights DeepHOPE as a potential non-invasive tool for guiding and timing the selection of nascent hiPSC colonies directly on culture plates during reprogramming processes.

**Figure. 4.**
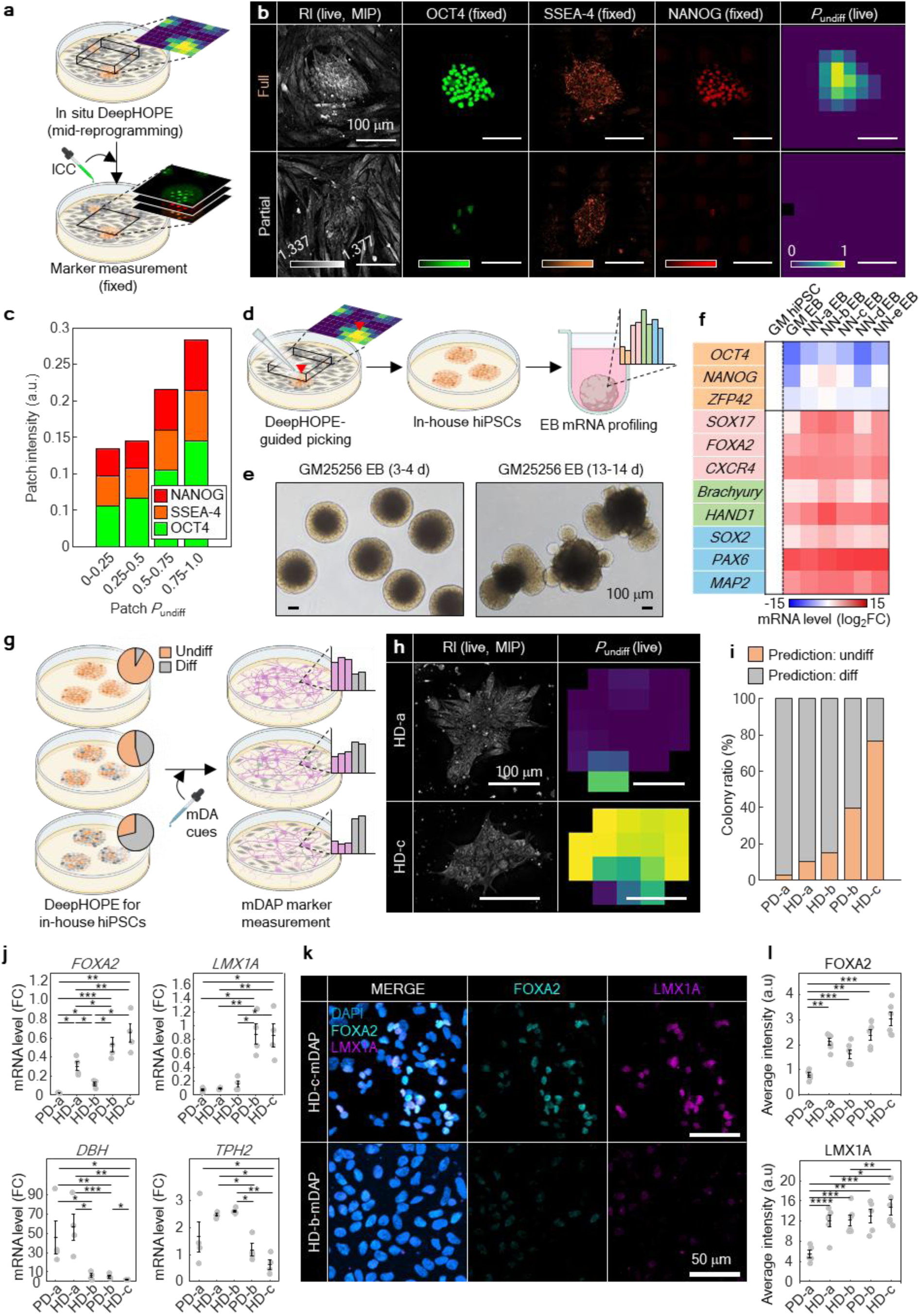
DeepHOPE-guided selection of functionally competent hiPSC colonies for personalised stem cell therapy. **(a)** Schematic of mid-reprogramming demonstration of DeepHOPE followed by marker-based validation. **(b)** Representative HT (live), fluorescence (fixed) images (OCT4, SSEA4, NANOG) and *P*_undiff_ maps at two locations of reprogramming culture, displaying distinct reprogramming outcomes. **(c)** Stacked bar plot of marker fluorescence intensities across ranges of patch *P*_undiff_ from reprogramming culture. *n* = 5 colonies. **(d)** Schematic of DeepHOPE-guided colony picking and followed by EB assay. **(e)** Microscope images of EBs formed from a standardised hiPSC line GM25256. **(f)** Measurements of pluripotency- and lineage-specific markers after EB formation. The two leftmost columns are measurements from GM25256 hiPSCs and EBs, providing the typical shift of marker expression. **(g)** Schematic of assessing DeepHOPE’s influence to the outcome of mDAP differentiation. **(h)** DeepHOPE-derived *P*_undiff_ of HD and PD patient-derived hiPSCs before differentiation (Day 0). **(i)** Rates of colony-level DeepHOPE prediction across multiple reprogrammed hiPSC batches. *n* = 15 (PD-a), 14 (HD-a), 12 (HD-b), 6 (PD-b) and 4 (HD-c) colonies. **(j)** Normalised mRNA expression levels of mDAPs markers at day 15. **(k)** Fluorescence images of LMX1A and FOXA2 in mDAPs derived from individual hiPSC lines at day 15. **(l)** Fluorescence intensity of LMX1A and FOXA2 in mDAPs at day 15. *n* = 4 independent mDAP differentiations [(J) – (L)]. Bar represents mean ± SEM and asterisks denote \**p* < 0.05, \*\**p* < 0.01, \*\*\**p* < 0.001 and \*\*\*\**p* < 0.0001 in Welch’s *t*-test between two adjacent or non-adjacent intervals.

To assess whether DeepHOPE-based evaluation during reprogramming reflects the later differentiation potential of selected cells, we performed embryoid body (EB) differentiation assays. Regions exhibiting high DeepHOPE-derived *P*_undiff_ within fibroblast reprogramming cultures were isolated and transferred into an EB-formation environment, consisting of suspension culture under spontaneous differentiation conditions (Fig. 4d,e). Gene expression profiling confirmed robust induction of lineage-specific markers representing all three germ layers—endoderm, mesoderm, and ectoderm—at levels comparable to those observed in EBs generated from the well-characterised GM25256 hiPSC line (Fig. 4f and Extended Data Fig. 7). These results demonstrate that DeepHOPE not only enables label-free pluripotency assessment during reprogramming but also reliably identifies colonies with strong trilineage differentiation potential for diverse downstream applications.

We further evaluated the impact of DeepHOPE-guided hiPSC selection on the efficiency of hiPSC differentiation into specific cell types—an important metric for personalised stem cell therapies. During the manufacturing of cell therapy products, transient fluctuations in stem cell quality can lead to substantial differences in both functional outcomes and safety profiles^43^. To demonstrate this, we compared the production of midbrain dopaminergic neuron progenitors (mDAPs), a key cell population for Parkinson’s disease cell therapy, among in-house hiPSC lines that differed in their DeepHOPE-derived assessments (Fig. 4g). The five hiPSC lines, reprogrammed from HD and PD fibroblasts, exhibited distinct ratio of colonies predicted to be “undifferentiated” by DeepHOPE (Fig. 4h,i). By day 15 of differentiation, the expression levels of mDAP markers, including LMX1A and FOXA2, aligned with the DeepHOPE predictions, indicating that DeepHOPE-based screening enhances the efficiency of generating targeted cell types (Fig. 4j-l). Furthermore, derivatives from hiPSC lines with high proportion of DeepHOPE-classified ‘differentiating’ colonies displayed elevated expression of non-mDAP markers, such as DBH⁺ (adrenergic) and TPH2⁺ (serotonergic) populations, which may cause adverse effects following clinical transplantation, including dyskinesia^44,45^.

### Colony flattening as a key structural indication of early exit from pluripotency

To elucidate the structural basis of DeepHOPE predictions, we profiled 31 topological, geometric, and compositional properties derived from the 3D RI distribution, including average RI, roundness, boundary contrast, average thickness, lipid volume, intercellular gap area, kurtosis, and centroidal thickness displacement (Extended Data Fig. 8 and 9b). Based solely on these properties, we clustered individual colonies and observed that each differentiation status occupied a characteristic region in the resulting property space (Fig. 5a and Extended Data Fig. 9a). To identify the features most strongly associated with DeepHOPE predictions, we overlaid colony-level *P*_undiff_ onto the feature map and ranked clusters by their average *P*_undiff_ (Fig. 5b). This analysis revealed that aspect ratio, curvature, and average thickness were major contributors to DeepHOPE predictions; notably, the cluster with the highest average *P*_undiff_ showed pronounced differences in these properties relative to other clusters (Fig. 5c,d). We further examined the major properties with the highest correlation between properties and *P*_undiff_ and identified that the average thickness showed strong correlation with *P*_undiff_ (Extended Data Fig. 9c). We next quantified topographical transformations of colonies during early differentiation by comparing colony surface profiles and tracking average thickness over time (Fig. 5e,f and Extended Data Fig. 10). Interestingly, we observed colony flattening across all early stages of differentiation up to 24 hours, whereas later time points exhibited more divergent patterns depending on the differentiation trajectory. Consistent with this observation, gradient-weighted class activation mapping (Grad-CAM) frequently highlighted flattened regions as contributing to classification into the “differentiating” state (Extended Data Fig. 11a-c). To further validate whether colonial flattening correlates with actual pluripotency protein expression, we applied the same analysis method on isolated fibroblast-derived hiPSC lines. Across fibroblast-derived hiPSC lines, each line occupied a broad range within the property space, underscoring substantial intra-batch structural heterogeneity (Fig. 5g). Nevertheless, colony flattening—reflected by a reduced aspect ratio and average thickness—was consistently prominent among colonies with low *P*_undiff_ and diminished pluripotency marker expression (Fig. 5h,i).

**Figure. 5.**
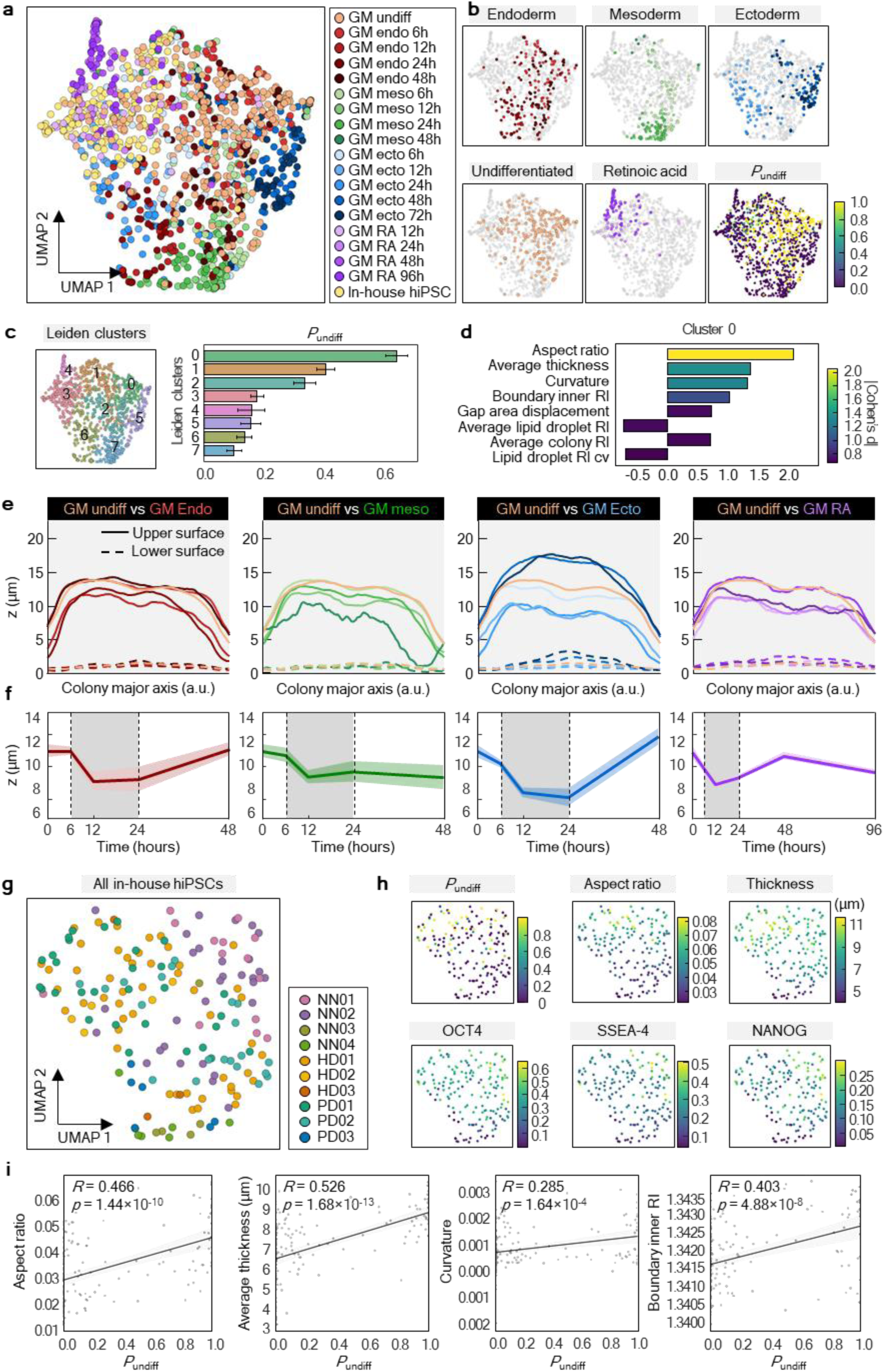
HT-derived colony properties related to DeepHOPE result and pluripotency marker shift. **(a)** UMAP representation of HT-derived quantitative properties of individual colonies including topological, geometric, and compositional properties. Colonial status and differentiation time-courses are colour-coded: GM25256-undifferentiated (white orange), endodermal differentiation (red colour gradients), mesodermal differentiation (green colour gradients), ectodermal differentiation (blue colour gradients), RA-induced differentiated (purple colour gradients), and reprogrammed hiPSCs (white yellow). **(b** and **c)** *leiden* clustering (*kNN* = 8) by morphological properties and the average colony-level *P*_undiff_ of each cluster. **(d)** Top properties contributing to the separation of cluster 0 from other clusters. **(e)** Average surface profiles along each colony’s major axis. The major axes of different lengths were all scaled to 0 to 1, with overall lowering surface high. **(f)** Colony thickness dynamics during three germ layer differentiation and RA-induced differentiation. Shaded time-courses between dotted bar remarks colony flattening. **(g)** UMAP clusters of reprogrammed hiPSCs from HD, PD and BJ fibroblasts. **(h)** Comparison of *P*_undiff_, aspect ratio, and curvature with pluripotency marker (OCT4, SSEA4, NANOG) expression. **(i)** Linear fit between *P*_undiff_ and aspect ratio, average thickness, curvature and average RI MIP at the inner boundary of each colony. *R* and *p* refer to Pearson’s correlation coefficient and *p*-value respectively.

### F-actin remodelling rapidly mediates colony flattening during early pluripotency exit

To elucidate the mechanism underlying rapid colony flattening during early differentiation, we examined cytoskeletal remodelling, with a particular focus on F-actin redistribution. Previously, F-actin was known to play a critical role in maintaining apical–basal polarity and holding colonial contractility in hPSCs^46–48^, and we hypothesised that the colony flattening was driven by F-actin redistribution. As expected, F-actin underwent immediate and pronounced redistribution within 24 hours of differentiation induction, with apical F-actin decreasing and basal F-actin increasing. (Fig. 6a). Compared to the undifferentiated state, differentiation toward all three germ layers showed reduced apical (z = 9.496 ∼ 11.395 μm) F-actin intensity and dry mass. In contrast, basal (z = 1.899 ∼ 3.798 μm) F-actin intensity and dry mass were enhanced (Fig. 6b,c). F-actin was consistently rearranged in parallel with changes in dry mass, and the bidirectional changes in apical and basal domains were collectively associated with colony flattening.

**Figure. 6.**
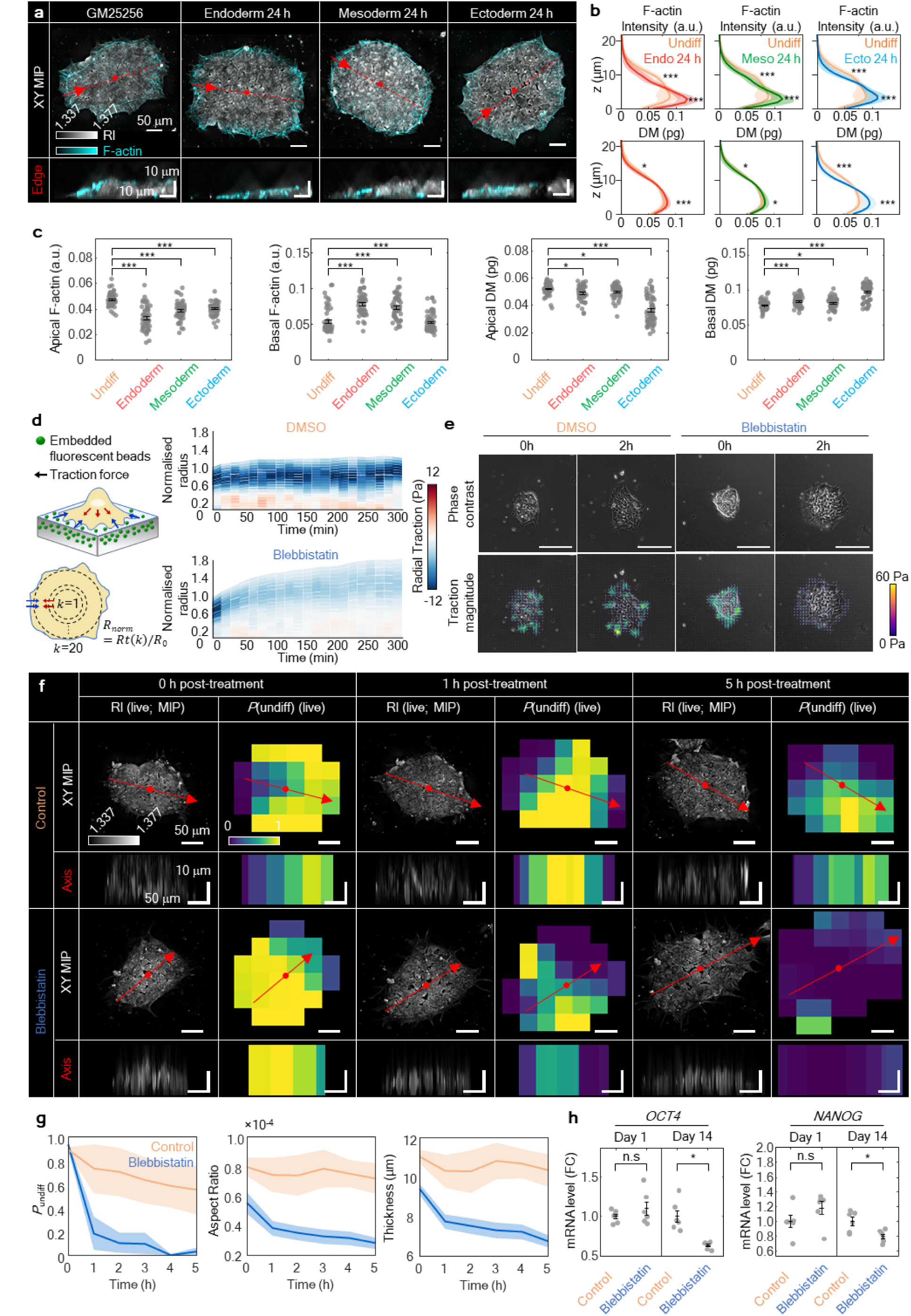
DeepHOPE-based detection of minute changes in colony architecture during early differentiation associated with F-actin remodelling. **(a)** Co-imaging results of HT and F-actin fluorescence. Scale bar = 50 μm (top) and 10 μm (bottom). **(b** and **c)** F-actin fluorescence intensity and dry mass distribution along the *z*-axis. *n* = 56 (Undifferentiated), 49 (Endoderm), 49 (Mesoderm) and 61 (Ectoderm) colonies. **(d)** Time-course radial traction force change after blebbistatin treatment in GM25256. Time interval = 15 minutes. *n* = 5 colonies for each group. **(e)** Representative images of blebbistatin-induced traction force changes. Scale bar = 100 μm. **(f)** Representative images of blebbistatin-induced morphological changes. Time-course of *P*_undiff_ evaluated by DeepHOPE. Each section view was generated at the edge region along colony major axis. Scale bar = 50 μm (top view) and 10 μm (section view). **(g)** Time-course of average *P*_undiff_, aspect ratio and thickness. *n* = 4 (DMSO) and 6 (blebbistatin). **(h)** mRNA expression of *OCT4* and *NANOG* during blebbistatin treatment at day 1 and day 14. *n* = 6 for each group. In [(B), (C) and (H)], Bar represents mean ± SEM and asterisks denote \**p* < 0.05, \*\**p* < 0.01, \*\*\**p* < 0.001 and \*\*\*\**p* < 0.0001 in Welch’s *t*-test.

To further examine whether F-actin-mediated topological transition functions as a major criterion in DeepHOPE, we treated colonies with low-dose blebbistatin to quickly reduce actomyosin tension^49,50^ and used TFM to check the immediate mechanical response^51,52^. Within 15 minutes, traction across the whole colony dropped, before we could see obvious spreading or flattening, suggesting that force loss happens first and may trigger the shape change (Fig. 6d). In DMSO controls, traction hotspots persisted, and an organised inward radial traction pattern appeared near the periphery, consistent with stable perimeter-associated contractility. In contrast, blebbistatin broadly weakened traction and disrupted this inward pattern while reducing edge counter-stress (Fig. 6e). DeepHOPE rapidly detected blebbistatin-induced morphological changes and displayed low *P*_undiff_ at the flattened regions (Fig. 6f). With colony flattening, aspect ratio, curvature and average thickness began to decrease within 1 hour after blebbistatin treatment, and the corresponded *P*_undiff_ rapidly declined (Fig. 6g). We further examined whether manipulation of F-actin alone is sufficient to influence pluripotency exit and found that long-term treatment of blebbistatin for 14 days induced a reduction in pluripotency (Fig. 6h). In summary, these results suggest that F-actin dynamics have a crucial role in maintaining pluripotency and respond sensitively to the initiation of differentiation.

## DISCUSSION

In this study, by integrating high-resolution HT with a spatially resolved DNN classifier, we developed DeepHOPE, a label-free, image-based framework for evaluating the pluripotency of live hPSC colonies. DeepHOPE enables quantitative, non-destructive assessment of colony-level stemness. The system partitions each colony into multiple regions and distinguishes local morphology based on subtle structural features indicative of early differentiation. While trained on a dataset derived from lineage-specific differentiation of a single hPSC line, DeepHOPE demonstrated robust performance across a broad spectrum of biological contexts—including RA-induced differentiation, in-house hiPSC lines, and nascent colonies emerging during somatic cell reprogramming. The strength of DeepHOPE arises from the synergy between HT and deep learning. HT provides high-resolution, quantitative, and reproducible 3D RI maps of live cells, generating data well-suited for extracting structural patterns through statistical learning. Notably, HT-derived structural properties revealed previously unrecognised 3D, topological, and compositional attributes of live hPSC colonies. This capability highlighted colony thickness as a critical morphological marker for pluripotency assessment, consistent with concurrent F-actin remodelling during early stages of differentiation. In contrast, a BF-based variant of the framework showed substantially lower predictive accuracy (Extended Data Fig. 6a-c), underscoring the unique value of HT data for pluripotency evaluation. Our data support that F-actin redistribution constitutes an early, necessary structural transition that accompanies and possibly precedes pluripotency exit, providing DeepHOPE an intermediary between molecular state and colony status. Also, our result on blebbistatin treatment shows that DeepHOPE detects changes prior to the decline of major mRNA expression, suggesting its sensitivity to early, pre-transcriptional state transitions.

In earlier studies, colony-scale morphological features such as flattening, edge effects, and mechanical responses in hPSCs have been extensively described in the context of patterning, crowding, and mechanotransduction, and are therefore not, by themselves, specific to pluripotency exit^53,54^. Prior approaches have largely relied on fluorescence-based, destructive assays, limiting their utility for continuous, in situ evaluation. In contrast, DeepHOPE provides a systematic, non-invasive, and predictive framework for assessing pluripotency of live hPSC colonies based on 3D structure. Notably, DeepHOPE shows potential sensitivity even at minute molecular marker shifts indicating that it captures early structural transitions rather than lineage commitment per se. Accordingly, rather than claiming that colony morphology defines pluripotency, we show that pluripotency exit is associated with reproducible patterns of structural transitions that are preserved across various contexts. DeepHOPE thus distinguishes these specific transitions from unrelated perturbations by identifying conserved signatures observed during bona fide pluripotency exit.

DeepHOPE nonetheless has limitations that point toward future technical refinements. First, the training dataset primarily comprised isolated colonies with diameters under 400 µm. In isolated colonies that grew larger, DeepHOPE-derived *P*_undiff_ decreased in the central region (Extended Movie 3), even in the absence of exogenous inducers. While previous reports also indicate flattening and pluripotency loss—biased to the neuroectodermal lineage—at the centre of large colonies^14,55,56^, expanding the dataset to accommodate a broader range of colony size would validate and improve DeepHOPE’s evaluation on larger colonies. Second, the current DeepHOPE workflow would benefit from improvements in computational efficiency to facilitate deployment in large-scale hPSC cultures. Advances in HT reconstruction speed, along with DNN training strategies, will help render DeepHOPE more suitable for high-throughput hPSC production and maintenance. On the other hand, the DNN inference of DeepHOPE is deployable on commodity hardware with 1.8 GB of GPU memory (see Materials and Methods), which is comparable to existing DNN–based vision workflows. This underlines that acceleration of HT reconstruction will further propel DeepHOPE towards accessibility in standard laboratory and manufacturing environments.

Despite its limitations, we demonstrated that DeepHOPE’s patch-based architecture enables spatially localised assessment of pluripotency, accommodating colonies of diverse shapes, sizes, and internal heterogeneity. This design allows the system to detect early, region-specific signs of differentiation that may otherwise be overlooked in global image analysis. When integrated with automated colony picking and culture systems, DeepHOPE has the potential to streamline the selective isolation of fully reprogrammed hiPSC colonies, thereby reducing unnecessary maintenance of suboptimal or non-functional lines^57,58^. Moreover, DeepHOPE-assisted mDAP differentiation enhanced robust and efficient production of stem cell therapy products for clinical application. Ultimately, DeepHOPE facilitates the scalable and cost-effective production of high-quality hiPSCs for personalised regenerative applications^59,60^.

In summary, DeepHOPE represents a significant advance in label-free pluripotency evaluation by integrating quantitative imaging with artificial intelligence (AI). Its broad applicability across diverse experimental settings, compatibility with live-cell workflows, and potential for integration into automated platforms highlight DeepHOPE as a versatile tool for stem cell quality control. As imaging and AI technologies continue to evolve^61–64^, DeepHOPE provides a blueprint for future systems that unify morphological, molecular, and functional insights in regenerative medicine.

## METHODS

### In vitro culture and maintenance of hPSCs

The human embryonic stem cell line H9 (WA09, WiCell) and human iPSCs lines GM25256 (Coriell Institute) and KOLF2.1J (The Jackson Laboratory) were used to generate the base model for this study. All hPSC lines were cultured on Matrigel (Corning, #354277)-coated plates with Essential 8 (E8) medium (Thermo Fisher, #A1517001). For passaging, cells were dissociated with 0.5 μM EDTA (Welgene, #LS015-10) and seeded onto fresh plates without ROCK inhibitor. Cells were cryopreserved using CryoStor® CS10 (STEMCELL Technologies, #07930) and stored in liquid nitrogen.

### Generation of human iPSCs from human dermal fibroblasts

To validate the model across various pluripotency levels of hiPSCs, three fibroblast samples (neonatal BJ, healthy donor-derived, and Parkinson’s disease patient-derived fibroblasts) were reprogrammed using mRNA-based reprogramming factors (Reprocell, #00-0076), containing OSKMLN (*Oct4, Sox2, Klf4, cMyc, Lin28* and *Nanog*). Reprogramming factors were transfected using Lipofectamine™ RNAiMAX Transfection Reagent (Thermo Fisher, #13778075). Infected fibroblasts were maintained in Nutristem (Sartorius, #05-100-1A) for 14-21 days, after which reprogrammed single colonies of hiPSCs were manually isolated and transferred onto Matrigel-coated plates in the presence of ROCK inhibitor (Y27632; Selleckchem, #S1049). Reprogrammed iPSCs were adapted and maintained in E8 medium for more than 10 passages prior to imaging and analysis. All reprogrammed hiPSCs were characterised by RT-qPCR to measure the expression of Oct4 and Nanog.

### Differentiation of hPSC into endoderm, mesoderm and ectoderm lineage

The GM25256 cell line (Coriell Institute) was primarily used for three germ layer-specific morphological analysis. Cells were dissociated with 0.5 μM EDTA (Welgene, #LS015-10) and seeded onto a fresh plate without ROCK inhibitor. Before inducing lineage-specific differentiation, cells were cultured in E8 medium for 24 hours, then lineage-specific differentiation was induced using STEMdiff™ Trilineage Differentiation Kit (STEMCELL Technology, #05230). The differentiation process was validated by RT-qPCR analysis on germ layer-specific markers.

### Differentiation of hPSC induced by retinoic acid

All-trans RA was purchased from Sigma-Aldrich (Sigma, #R2625). The retinoic acid powder was dissolved in dimethyl sulfoxide (DMSO; Sigma, #D8418) to prepare a stock concentration of 100 µM. To stably induce a low pluripotency status while maintaining an adequate colony size, 100 nM of retinoic acid was treated at 24 hours after replating 3 hPSCs lines (The time point of 0 hr), and cells were imaged up to 96 hours after replating. For the 12-hour sample, retinoic acid was treated once at 12 hours before imaging. For the 24-hour sample, to avoid the effects of retinoic acid half-life, retinoic acid was treated at 24 hours and 12 hours before imaging. For the 48-hour sample, retinoic acid was delivered at 48 hours, 24 hours, and 12 hours before imaging.

### Low-coherence holotomography

The 3D RI distribution of hPSCs was measured by a commercial low-coherence HT system (HT-X1, Tomocube, Daejeon, South Korea). At each focal depth, the LED illumination system (centre wavelength = 450 nm) illuminates the specimen with four distinct aperture profiles, which had been optimised to provide a uniform optical transfer function^65^. A digital micromirror device at the aperture plane rapidly switches between aperture patterns, while a custom-designed condenser lens (NA = 0.72, working distance = 30 mm) focuses the light onto the specimen. The diffracted light is collected by a 40× objective lens (Olympus, UPLXAPO40X, NA = 0.95, working distance = 0.18 mm). The objective lens is vertically translated with a motor to axially scan through focal depths. The stack of intensity measurement with varying illumination patterns and at a range of axial depths, are transformed into a 3D RI image through deconvolution based on the known optical transfer function. The motorised specimen stage allows horizontal movement of the field of view (FOV) and is equipped with an incubation chamber (TOKAI HIT, STXG-WSKMXA22B-E) to maintain the temperature, humidity, and CO2 level. The maximum FOV at a single stage coordinate is 165 µm × 165 µm, while the depth range of acquisition is 140 µm.

### Data acquisition

For live hPSC imaging, cells were cultured in E8 medium for at least three passages after thawing. One day before imaging, confluent cells were dissociated using 0.5 μM EDTA, and 30,000 – 50,000 cells were seeded onto a Matrigel-coated TomoDish (Tomocube, Daejeon, South Korea; #901002) without ROCK inhibitor. Cultures were maintained for 24–48 h prior to imaging. Horizontal image stitching, automated in TomoStudioX, was utilised to acquire a 3D RI tomogram equivalent to the size of a single hPSC colony. Specifically, image acquisition and reconstruction were performed individually at each horizontal coordinate, and the resulting single-FOV tomograms were then stitched together using relative coordinates that provide the highest correlation in the overlapping regions. As colony size ranged from 300 µm × 300 µm to 600 µm × 600 µm FOVs, the data acquisition for a single colony typically took 1 to 3 minutes.

### Data processing

Each HT image of an hPSC colony was numerically split into patches for lowered computational overhead. Each HT image was initially cropped along the vertical (*z*) axis to exclude regions unoccupied by cells, such as below the culture substrate surface. Then the horizontal 2D binary mask for the colony was determined first by image processing that includes RI thresholding (above 1.342) in MIP along the *z* direction, followed by two serial closing operations of circular kernels of 1-μm and 2-μm radii. Between the dilation and erosion of each closing operation, all enclosed background regions were filled. Followingly, each HT image was split into patches, within the square region that contains the binary mask with 20-μm wide margins in four directions if available. Each patch was of 119.3 μm × 119.3 μm size in case of the training set and 79.6 μm × 79.6 μm size for the validation set and the blind test set. The spacing between adjacent patches is 39.8 μm, in either *x* or *y* direction. The patch size and stride were selected to capture mesoscopic colony architecture while maintaining sufficient local context and computational tractability, and colony-level decisions were obtained by aggregating predictions across overlapping patches. The colony masking and patch creation were performed using MATLAB R2024a (MathWorks, Natick, MA, USA).

Additional on-line processing was carried out when feeding the stored patches into the DNN for training and inference. During training, a patch was cropped into 79.6 μm × 79.6 μm at a random coordinate in the horizontal direction, while cropped vertically within 11.4 μm above the substrate surface. The RI values were then rescaled to a range from 0 to 1, linearly mapped from the actual range between 1.330 and 1.400. Also, random alterations including horizontal flips (independent in *x* and *y* direction), *x*-*y* permutation, and noise addition were carried out for data augmentation. The noise was independent and identically distributed throughout all pixels, sampled from a Gaussian distribution at a standard deviation between 0.001 and 0.1 (randomly set every time). The flips and *x*-*y* permutation took place with a 50% probability. Finally, the vertical dimension was permuted to occupy the channel dimension of the DNN input. During inference, the random cropping was replaced with central cropping of identical size, while augmentation processes were removed. The on-line processing was realised mainly using NumPy 1.21.2 and PyTorch 1.10.0.

### DNN design and optimisation

The DNN design consists of a modified ResNet^66^ backbone enhanced with bottleneck attention modules (BAMs)^67^. After initial downsampling via strided convolution and pooling, the input image passes through four stages that progressively reduce the spatial size of the feature maps while increasing the feature map number. Finally, the network produces two logits—representing *undifferentiated* and *differentiating*—through global average pooling followed by a fully connected layer. The output feature map dimensions (channel × height × width) for the four stages are 256×64×64, 512×32×32, 1024×16×16, and 2048×8×8. Between neighbouring stages, a BAM is placed to aid the DNN focus on information localised in certain locations or channels. The four stages consist of three, four, six, and three bottleneck residual blocks, each of which is the smallest unit of residual image processing in ResNet. The first block of each stage reduces the feature map size by introducing 2×2 strides in both the convolutional path and residual path. Most of the convolutional layers in the DNN are followed by batch normalisation^68^ and rectified linear unit activation^69^. The DNN includes 23,981,749 parameters that mainly account for a total of 65 convolutional layers.

The optimisation was performed by gradient-based modification of DNN parameters, reducing the cross-entropy loss between the softmaxed DNN output and the one-hot ground truth vector. More specifically, the stochastic gradient descent (SGD) algorithm was employed at the batch size of 256, learning rate of 0.001, Nesterov momentum^70^ of 0.9, and weight decay of 0.0001. One iteration over the data reflected a randomly sampled 5% of the total training set data, to allow variability in the training. Also, the learning rate was modified after each iteration, following the cosine annealing method^71^ at the period of 16 iterations.

The graphics memory requirement for our DNN implementation was 87,314 MB and 1,810 MB in case of optimisation and inference respectively. We utilised a central processing unit of Xeon Silver 4114 (Intel Corporation, Santa Clara, California, USA) and four graphics processing units of Tesla P40 (Intel Corporation, Santa Clara, California, USA). The iterative optimisation consumed 53 minutes per iteration, and was carried out for a total of 300 iterations. The best-performing parameter checkpoint (264^th^ iteration in our case) was selected according to the highest sum of the validation accuracy and the blind test accuracy. The inference of an individual patch consumed 0.48 s under the inference batch size of 1.

### Saliency map visualisation

We employed Gradient-weighted Class Activation Mapping (Grad-CAM)^72^ to track the locations in the colony that the DNN recognises as signs of the undifferentiated or differentiated status. Grad-CAM is a class-discriminative saliency mapping, computed by a weighted sum of feature maps at a certain layer. The weight of each feature map is the gradient of the class output in the feature map space, averaged over all feature map coordinates. We obtained Grad-CAM maps using the feature maps from second convolutional layer of the last residual block.

### Deriving quantitative hPSC colony properties from HT measurement

Quantitative hPSC colony properties were extracted from each HT image, using rule-based processes. For the extraction of various properties via the 3D resolving power of HT, additional image processing was carried out to obtain 3D or 2D masks of the colony and specific structures such as intercellular gaps. First, the 3D mask of colony was obtained by taking the intersection between an RI-thresholded (RI > 1.3400) 3D region and the 2D mask of the colony in every *z* section. The 3D mask of colony lead to the calculation of colony volume, as well as the average and standard deviation of colony RI. Also, thickness profiling was available with the 3D mask, providing the average and standard deviation of thickness over the colony area. The upper surface of the colony was also used to calculate the curvature, by fitting the surface coordinates to a spherical surface. The thickness profile was considered within 5 μm or more inwards from the colony’s 2D boundary, to reduce bias from colony size or shape. The 3D location of intracellular lipid was further specified by thresholding the 3D HT image with a different range of RI. More specifically, lipid-occupied region was identified with high-RI (above 1.3800) region inside the colony, excluding those wider than 3 μm, unless the RI was below 1.4000, which are less likely to be lipid droplets in the perspective of size^73^. Another set of properties were measured to represent the 2D collective shape of each colony. For instance, roundness was calculated as the area ratio between the 2D colony mask and a sphere with the boundary length equal to the mask. The solidity was calculated as the area ratio between the 2D colony mask and the convex hull generated from the mask. Also, the eccentricity of the colony was defined as the eccentricity of the ellipse that fits the colony mask the most. Other properties are obtained by combining 3D and 2D morphologies as well as the RI distribution. The boundary RI contrast for each colony was also quantified as the difference between RI values averaged over the area 2 μm inwards and outwards the colony boundary. The area ratio of intercellular gaps was also assessable by determining the gap-occupied regions. The horizontal map for gaps was obtained as low-RI (below or equal to 1.3400) region in the horizontal MIP of the HT image. A gap region was excluded when 25% or more area was within the 5-μm proximity of the colony boundary, to prevent indented boundaries from being misdetected as gaps. Finally, the values that can be mapped onto 2D coordinates provide additional spatial properties including spread, skewness, and kurtosis. Applicable values include thickness, dry mass (obtained from RI), presence of lipid, and presence of gaps. HT images that resulted in the 2D colony masks whose bounding boxes were smaller than 14,250 μm^2^ in area were excluded from the property extraction—this mainly indicated a severe disruption of colony structure. All property calculations were carried out using MATLAB R2024a.

### hPSC colony properties-based morphometric analysis

Calculated morphological properties of all the single colony data were analysed for clustering. With 31 properties, colonies were clusterised using the Leiden algorithm on k-nearest neighbour graph constructed from standardised structural properties. UMAP was used for two-dimensional visualisation of the same properties space. For identification of top features contributing for each cluster, Cohen’s d of all the properties was calculated between one clusters and others. Morphometric calculations were carried out using Python 3.8.8.

### Quantitative reverse transcription polymerase chain reaction (qRT-qPCR)

Total RNA from hPSCs was extracted using the Ribospin™ II kit (#314-103, GeneAll) according to the manufacturer’s protocol. A total of 500 ng of RNA per 10 <u>μ</u>l of final reaction volume was used for reverse transcription using PrimeScript™ RT Master Mix (RR036B, Takara). For quantitative real-time polymerase chain reaction (qRT-PCR) analysis, cDNA was diluted 10-fold, and 5 <u>μ</u>l of diluted cDNA was mixed with GoTaq® qPCR Master Mix (Promega, A6101) according to the manufacturer’s protocol. qRT-PCR was performed using CFX Opus 96 Real-Time PCR System (BIO-RAD, #12011319). The sequences of qPCR primers are listed in Supplementary Table 2.

### Immunocytochemistry of hPSC colonies

To measure the expression level of pluripotency markers at the colony scale, hPSCs were labeled with antibodies against pluripotency markers: OCT4 (abcam, #ab19857), NANOG (R&D Systems, #AF1997, and SSEA4 (Millipore, #MAB4304). After live-HT imaging, hPSCs were immediately fixed with 4% paraformaldehyde (PFA) for 10 min at room temperature. The fixed cells were then blocked with a blocking buffer (5% donkey serum, 3% BSA, and 0.2% Triton X-100 in DPBS) for 30 min at room temperature. The primary antibody was diluted in the blocking buffer and added to the blocked cells. The cells were incubated with the primary antibody for 4 hours at room temperature, followed by three washes with PBST (0.1% Tween 20 in DPBS). The secondary antibody diluted in the blocking buffer was added and incubated in the dark for 1 hour at room temperature. After incubation, cells were washed three times with PBST and filled with DPBS for fluorescence imaging. Fluorescence imaging was performed using epifluorescence microscopy integrated into the HT-X1 system.

### Embryoid body formation and differentiation potential assay

To validate the differentiation potential of DeepHOPE-selected hiPSC colonies during reprogramming, individually iPSC clones were isolated and formed into embryoid body (EB). Cells were dissociated with 1X Accutase solution (#A6964-500ML, Sigma-Aldrich) and resuspended with Essential 8 medium containing ROCK inhibitor. Approximately 2,000 cells were evenly seeded onto 96-well Clear Round Bottom Ultra-Low Attachment Microplate (#7007, Corning) and centrifuged at 200 g for 5 min. Aggregated cells were incubated for 1 day and the next day, medium was changed to Essential 6 media (#A1516401, Thermo Fisher). 14 days after withdrawal of TGFβ and bFGF, differentiated EBs were lysed for RNA extraction using Ribospin™ II kit (#314-103, GeneAll) according to manufacturer’s protocol. Differentiation potential was evaluated by measuring early three-germ layer marker expression using RT-qPCR.

### Preparation of polyacrylamide gel substrates

Polyacrylamide (PA) gel substrates with a Young’s modulus of 3 kPa were prepared by mixing acrylamide and bis-acrylamide solutions according to predefined formulations^74,75^. For the 3 kPa gels, the PA solution consisted of 2042 μL deionised water, 344 μL of 40% (w/v) acrylamide solution (#1610140, Bio-Rad), and 112.5 μL of 2% (w/v) N,N′-methylenebisacrylamide solution (#1610142, Bio-Rad). Fluorescent beads (12.5 μL; diameter = 0.5 μm; FluoSpheres; #F8812, Invitrogen) were added prior to polymerisation. Polymerisation was initiated by adding 0.05% (v/v) ammonium persulfate (APS; #A3678, Sigma-Aldrich) and 0.05% (v/v) N,N,N′,N′-tetramethylethylenediamine (TEMED; #T9281, Sigma-Aldrich). A 24-μL drop of the PA solution was deposited onto a silane-treated glass surface and covered with a coverslip to control gel thickness. After polymerisation, the coverslip was removed, and the glass substrate was centrifuged to localise the fluorescent beads near the surface of the PA gel. For surface functionalisation, the PA gel was treated with sulfosuccinimidyl-6-(4-azido-2-nitrophenyl amino) hexanoate (Sulfo-SANPAH; #C1111, Proteochem) dissolved in 50 mM HEPES buffer (#H0887, Sigma-Aldrich) at a concentration of 1 mg/mL, followed by coating with basement membrane matrix.

### Time-lapse traction force microscopy

Time-lapse traction force microscopy (TFM) experiments were performed on an inverted microscope (Zeiss, Axio Observer series) equipped with a climate-controlled chamber (37 °C and 5% CO₂)^76,77^. Phase-contrast and fluorescence images were acquired every 15 min for approximately 6 h using a 10× objective lens. GM25256 cells were seeded onto the PA gels to allow colony formation. After 24 h of culture, time-lapse imaging was performed and traction maps were reconstructed from bead displacement fields. The projected area of the cell colonies was discretised into quadrilateral elements, and the x- and y-components of traction were calculated at each mesh node based on local bead displacements. Following imaging, the medium was replaced with blebbistatin-containing medium (final concentration = 5 μM; #B0560, Sigma-Aldrich) dissolved in DMSO to relax cytoskeletal tension and obtain the reference images. DMSO-treated samples served as vehicle controls.

## Corresponding authors

Further requests for data should be directed to and will be fulfilled by the lead contact, Ki-Jun Yoon.

## Code availability

The source code for the DeepHOPE framework is available at the GitHub repository (https://github.com/GeonKim94/DeepHOPE). Trained DNN checkpoints and sample data are available from the corresponding author upon reasonable request.

## Ethical Statement

The collection of fibroblasts was conducted following the approval of the Institutional Review Board (IRB) of the Research Ethics Committee at Seoul National University Hospital, Seoul, South Korea (IRB no.H-0905-041-281) and of the Public Institutional Review Board (IRB no.2021-3687-012). Written informed consent was obtained from all donors prior to biopsy.

## ACKNOWLEDGMENT

We thank all the members of the Park laboratory and the Yoon laboratory for their valuable comments and discussions, Jennifer H. Shin for thoughtful feedback and insightful discussions, Tomocube, Inc. for technical support on HT-X1^TM^, Bokyung Yoo and Byeongsuk Kim for their technical assistance. This work was supported by National Research Foundation of Korea (NRF) grants (RS-2023-00241278 and RS-2024-00332454 to G.K., RS-2024-00440778 and RS-2024-00332454 to K.-J.Y., RS-2024-00442348RS-2024-00440577 to Y.K.P.) funded by Korea Institute for Advancement of Technology (KIAT) through the International Cooperative R&D program (P0028463), the Korean Ministry of Science, ICT, and Future Planning (MSIP), and the Young Investigator Grant from the Suh Kyungbae Foundation (to K.-J.Y.), The graphical abstract was created with BioRender.com.

## AUTHORS CONTRIBUTION

H.P., G.K., Y.K.P. and K.-J.Y. contributed to the overall study design and conceptualisation. H.W. and G.K. performed all analysis and wrote the draft, and Y.K.P. and K.-J.Y. reviewed and edited the original draft. H.P., J.S., G.Y. and T.H. contributed to hPSCs culture, differentiation and HT imaging. G.K. contributed to deep learning model generation and structural properties extraction. S.-H.K., E.-B.H., M.K. and J.P. contributed to collecting donor-driven fibroblast and reprogramming. Y.J. contributed to measurement of traction forces and analysis. J.H.S., N.-S.K., Y.K.P. and K.-J.Y. supervised and provided critical feedback to results and method development.

---

**Extended Data Fig. 1.**
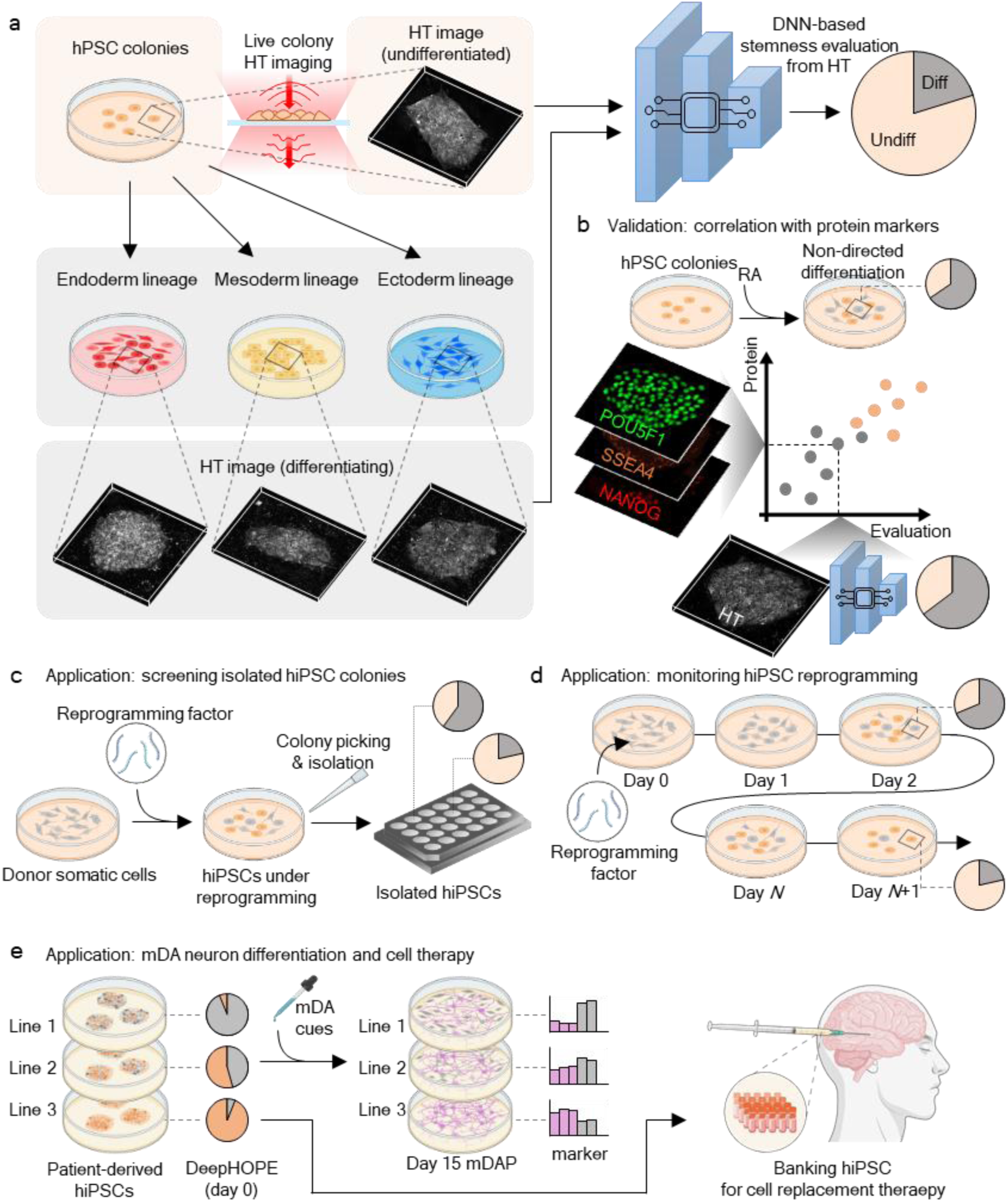
Overview of the DeepHOPE framework for label-free pluripotency evaluation. **(a**) Schematic of DeepHOPE workflow. Live human pluripotent stem cell (hPSC) colonies are imaged using holotomography (HT) to reconstruct three-dimensional refractive index (RI) maps without labelling. To train a deep neural network (DNN), HT images of undifferentiated and differentiating hPSC colonies are acquired under directed differentiation conditions towards endoderm, mesoderm, and ectoderm lineages. (**b**) The DNN is optimised to classify image patches from hPSC colonies, before and after RA treatment, into undifferentiated or differentiating categories, enabling spatially resolved inference of pluripotency status across the entire colony. The prediction outputs are benchmarked against immunostaining for key pluripotency markers, including POU5F1, SSEA4, and NANOG. (**c**) Applications of DeepHOPE include non-invasive screening of isolated hiPSC colonies following reprogramming and *in situ* monitoring of pluripotency dynamics during reprogramming. (**d**) The framework enables real-time, label-free assessment of colony-level stemness across diverse experimental contexts. (**e**) Clinical application of the framework for qualification of personalized hiPSCs before differentiation and the generation of cell therapy products.

**Extended Data Fig. 2.**
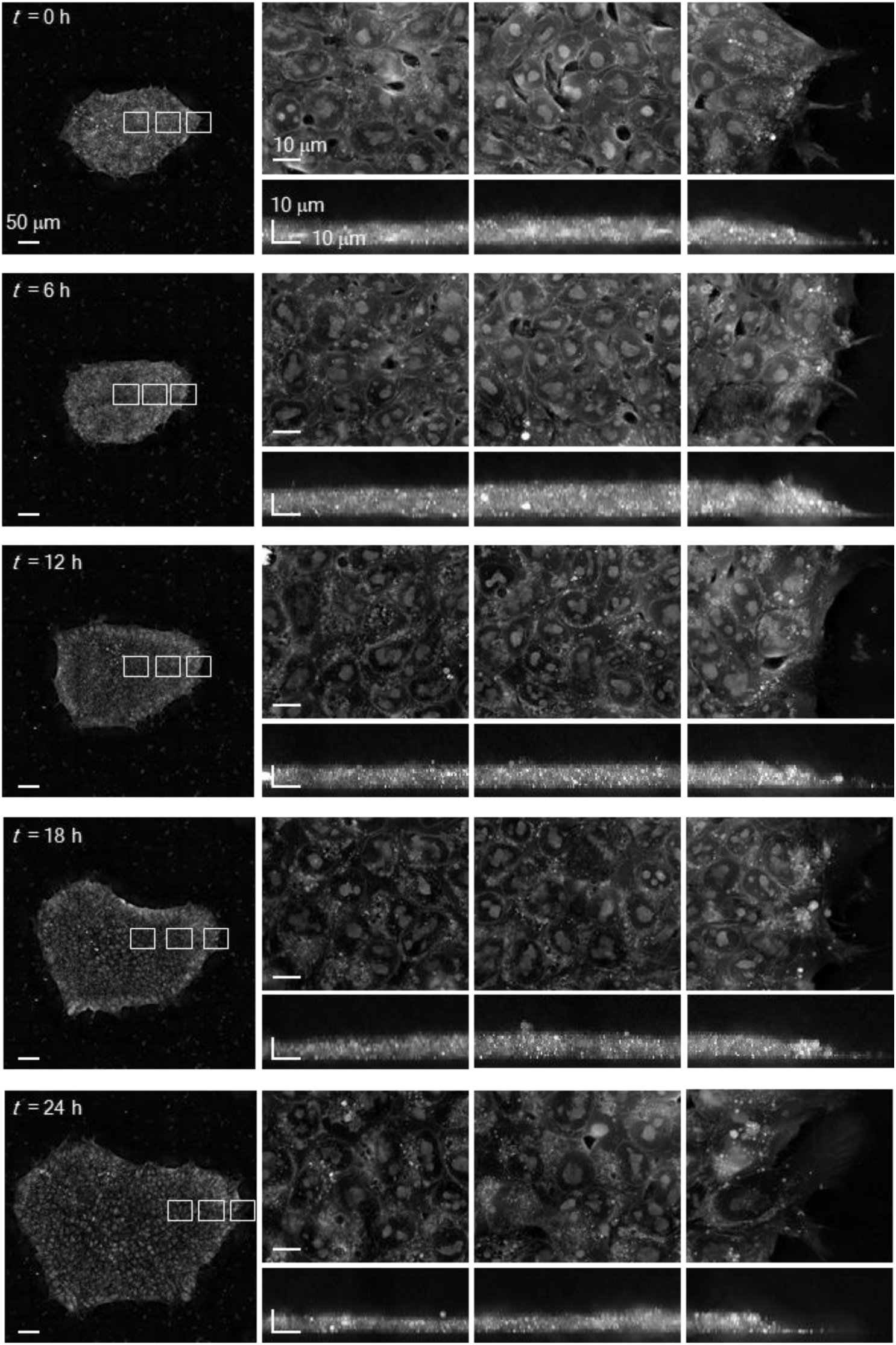
Time-lapse HT imaging of an hiPSC colony. Time-lapse images of a GM25256 colony over time. The right panels show regions indicated by insets in the leftmost images. All images are maximum intensity projections along the vertical (top row) or horizontal (bottom row) axis. The initial time point is 48 h post-subculture. Scale bar = 50 μm (left) and 10 μm (right).

**Extended Data Fig. 3.**
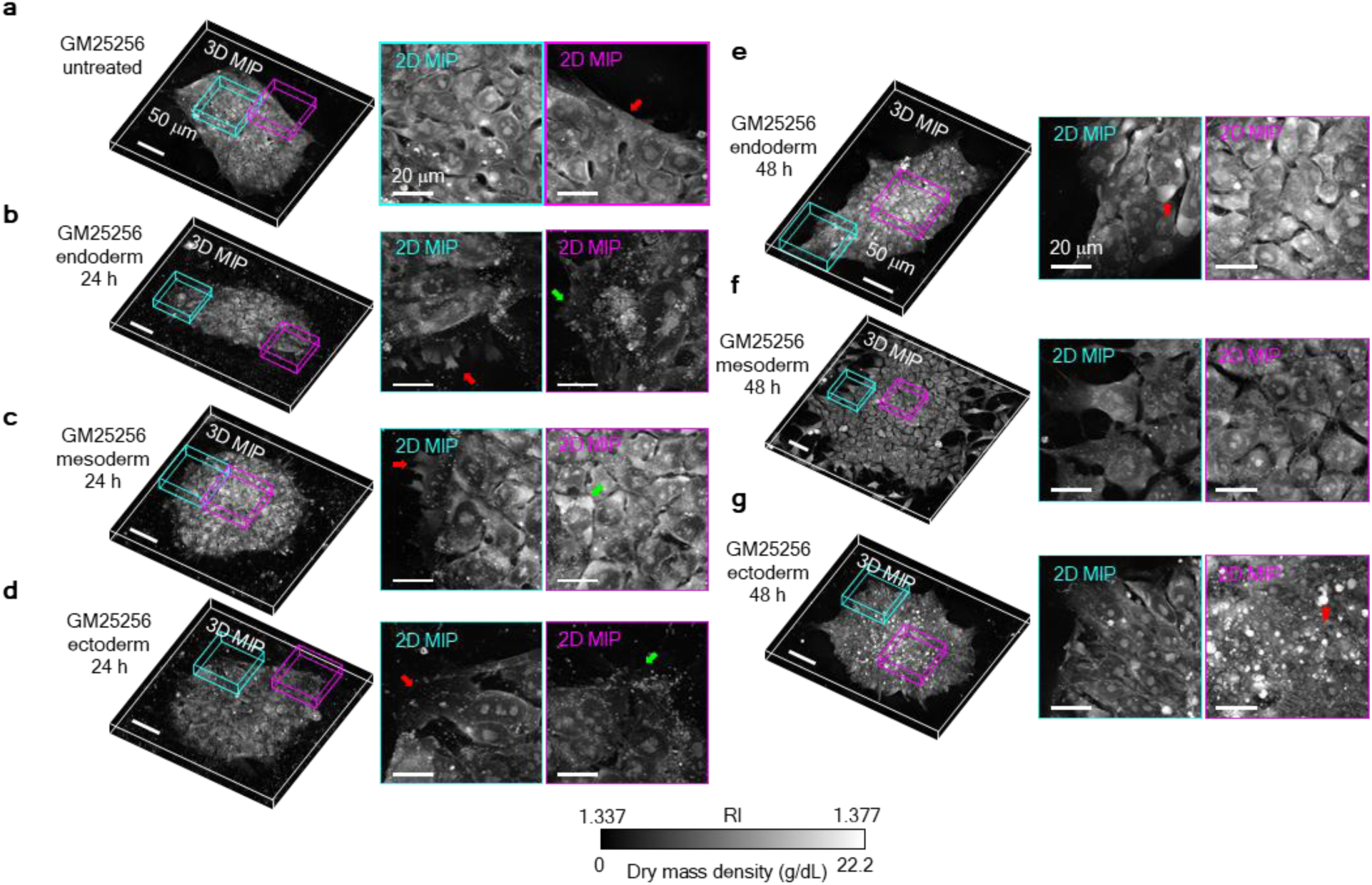
HT-measurable morphologies related to differentiation into three germ layers. **(a)** A GM25256 hiPSC colony under no induction of differentiation. The red arrow indicates a smooth and high-contrast boundary of the colony. **(b)** A GM25256 hiPSC colony after 24 h of endodermal differentiation. The red arrow indicates membrane protrusion at colony periphery. The green arrow indicates filopodium-like spikes at colony periphery. **(c)** A GM25256 hiPSC colony after 24 h of mesodermal differentiation. The red arrow indicates membrane protrusion at colony periphery. The green arrow indicates high-RI intracellular blobs. **(d)** A GM25256 hiPSC colony after 24 h of ectodermal differentiation. The red arrow indicates obscure colony boundary. The green arrow indicates filopodium-like spikes at colony periphery. **(e)** A GM25256 hiPSC colony after 48 h of endodermal differentiation. The red arrow indicates high-RI intracellular blobs. **(f)** A GM25256 hiPSC colony after 48 h of mesodermal differentiation. **(g)** A GM25256 hiPSC colony after 48 h of ectodermal differentiation. The red arrow indicates high-RI bodies among tightly-packed cells. Scale bar (**a** to **g**) = 50 μm (left) and 20 μm (right).

**Extended Data Fig. 4.**
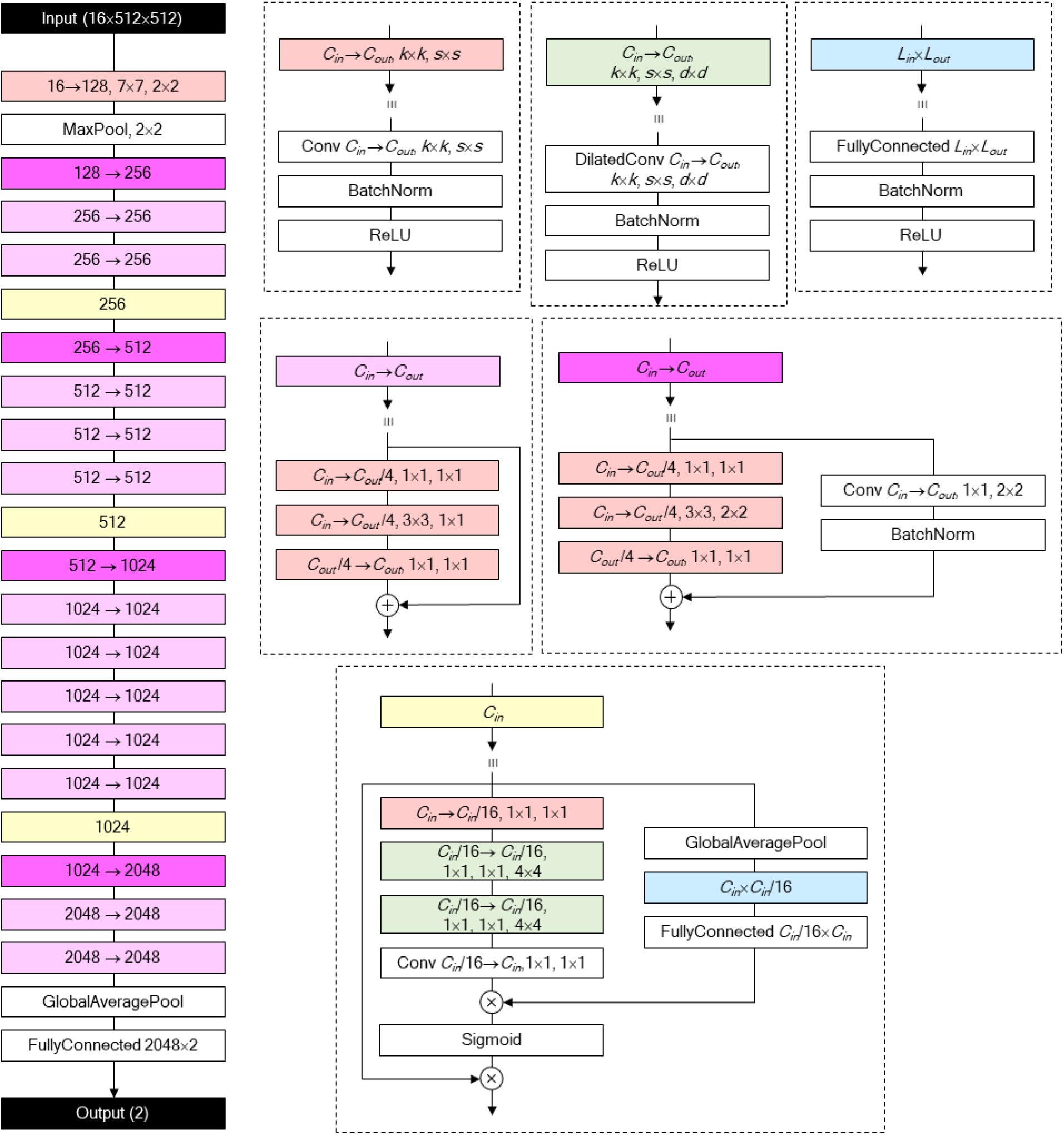
Structure of DNN for DeepHOPE evaluation. Each box refers to a computational operation. Boxes without colours denote basic operations while each of those with colours is composed of multiple basic operations (defined on the right). *C_in_*: input channel size of a convolution layer. *C_out_*: output channel size of a convolution layer. *K*: kernel size. *S*: stride size. *D*: dilation size. *L_in_*: input vector size of a fully-connected layer. *L_out_*: output vector size of a fully-connected layer. ⊕: concatenation. ⊗: Hadamard product.

**Extended Data Fig. 5.**
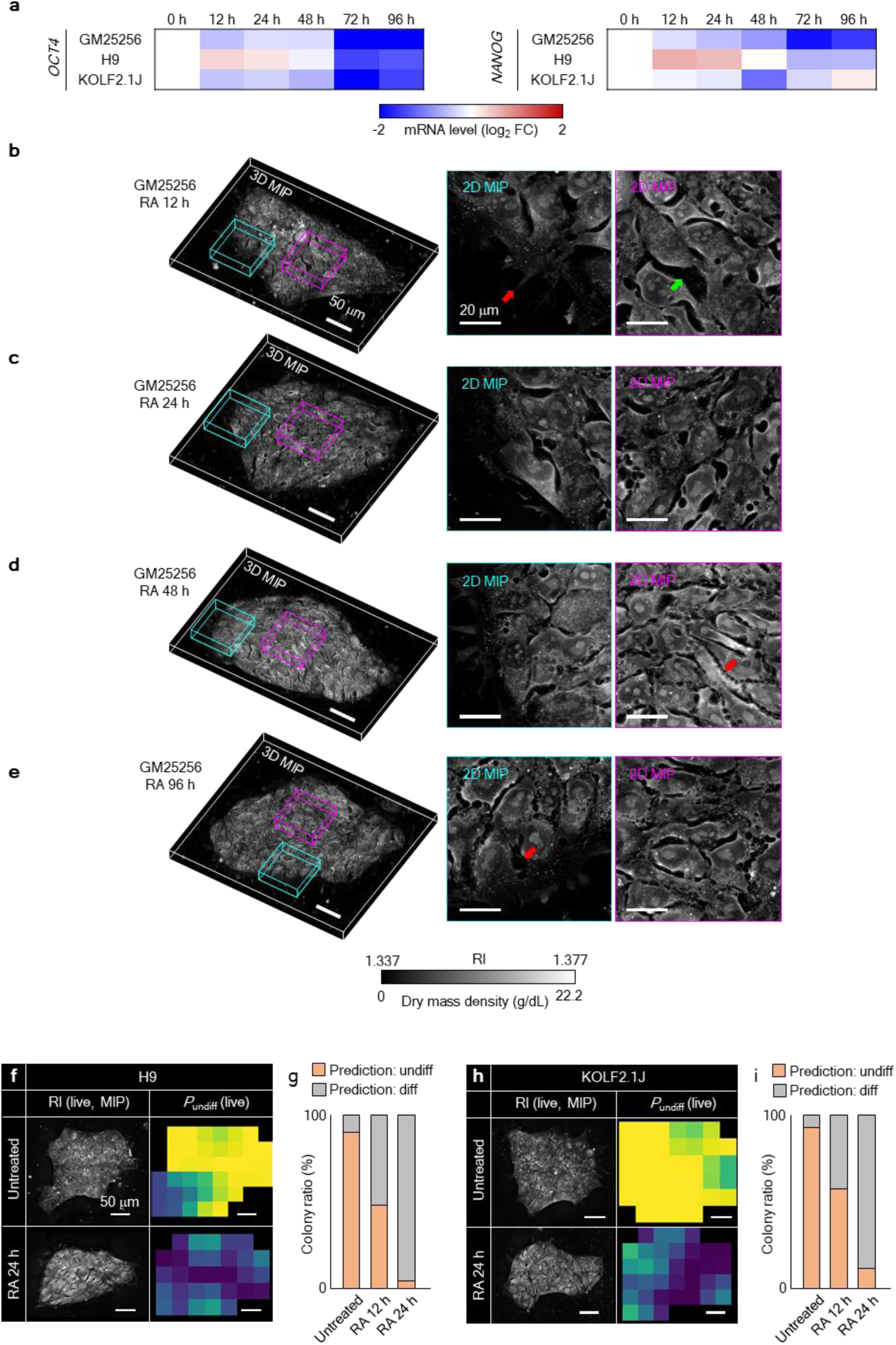
HT-detectable morphologies associated with RA-induced differentiation. **(a)** Transcriptomic analysis of the time-course differentiation process. Loss of pluripotency markers is visible over time. **(b)** A GM25256 hiPSC colony after 12 h of RA-induced differentiation. The red arrow indicates filopodium-like membrane protrusion at colony periphery. The green arrow indicates an intercellular gap. **(c)** A GM25256 hiPSC colony after 24 h of RA-induced differentiation. **(d)** A GM25256 hiPSC colony after 48 h of RA-induced differentiation. The red arrow indicates a jagged colony boundary. (**e**) A GM25256 hiPSC colony after 96 h of RA-induced differentiation. The red arrow indicates an intercellular gap near the colony periphery. (**f**) A H9 hESC colony both untreated and after 24 h of RA-induced differentiation. *P*_undiff_ was measured by DeepHOPE. **(g)** Ratio of H9 hESC colonies with average *P*_undiff_ over 0.5. **(h)** A KOLF2.1J hiPSC colony both untreated and after 24 h of RA-induced differentiation. *P*_undiff_ was measured by DeepHOPE. **(i)** Ratio of KOLF2.1J hiPSC colonies with average *P*_undiff_ over 0.5. Scale bar (B to H) = 50 μm.

**Extended Data Fig. 6.**
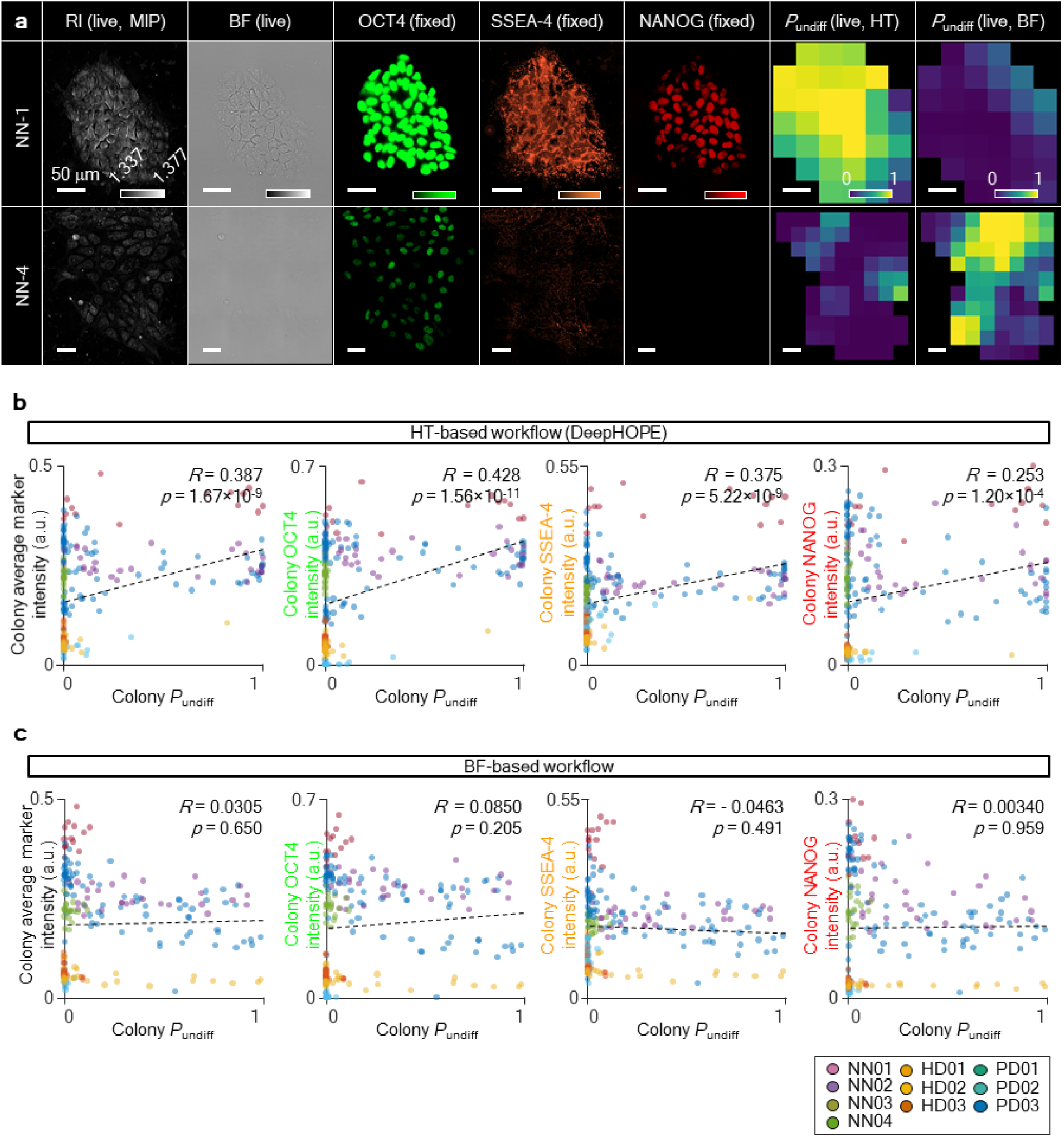
Comparison of *P*_undiff_-marker correlation between DeepHOPE and a bright field image-based alternative. **(a)** Representative HT image (left) and bright field (BF) image (right) of the same live GM25256 colony. Scale bar = 50 μm. **(b)** Linear fit between DeepHOPE *P*_undiff_ and marker fluorescence intensities in individual in-house hiPSC colonies. **(c)** Linear fit between BF-based *P*_undiff_ and marker fluorescence intensity of individual in-house hiPSC colonies. *R* and *p* refer to Pearson’s correlation coefficient and *p*-value, respectively.

**Extended Data Fig. 7.**
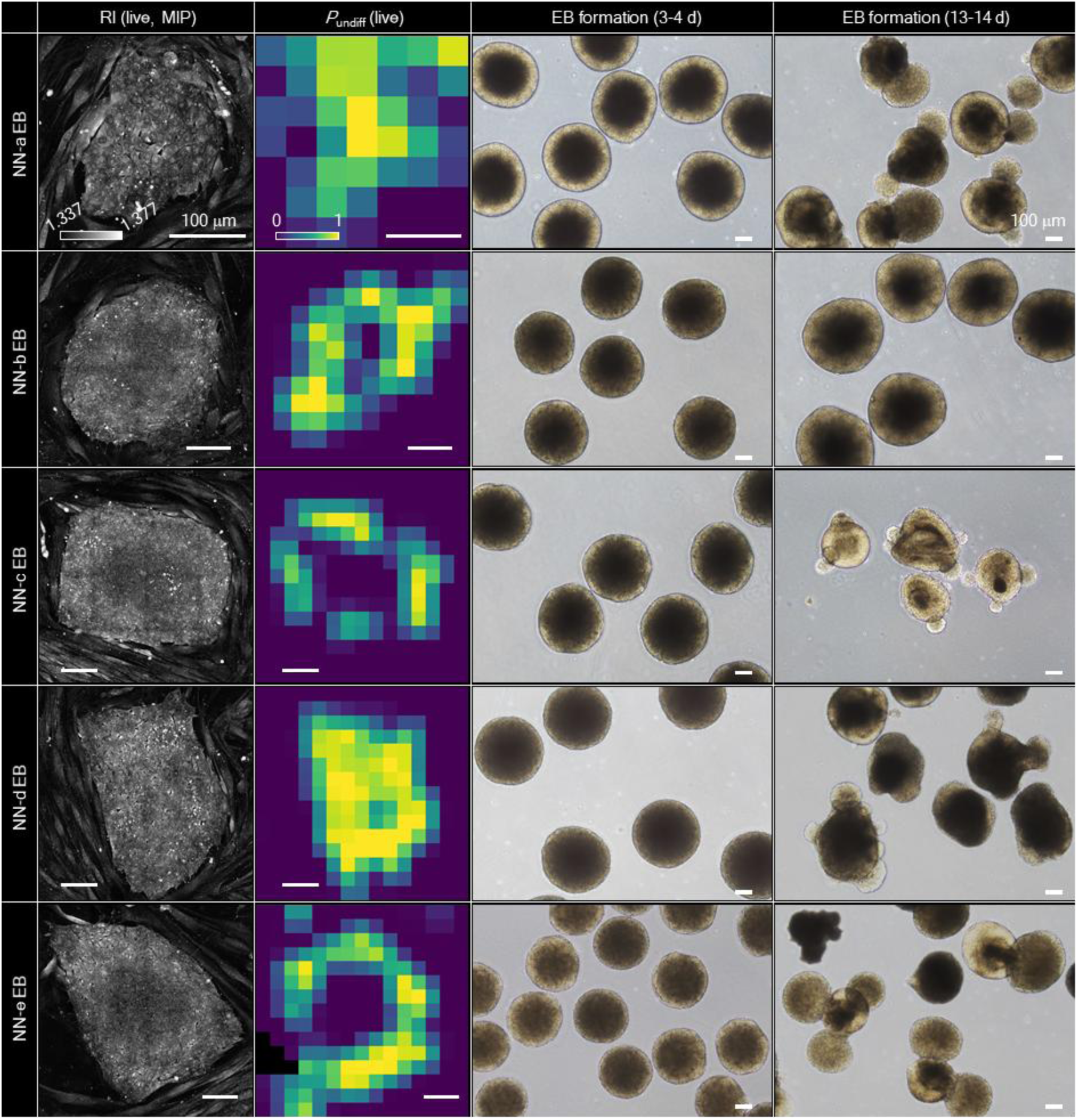
EB assay following DeepHOPE-guided colony picking. Each column shows (from left to right): the live HT image, the *P*_undiff_ map provided by DeepHOPE, the BF image of EBs cultured for 3–4 days, and the BF image of EBs cultured for 13–14 days. Scale bar = 100 μm.

**Extended Data Fig. 8.**
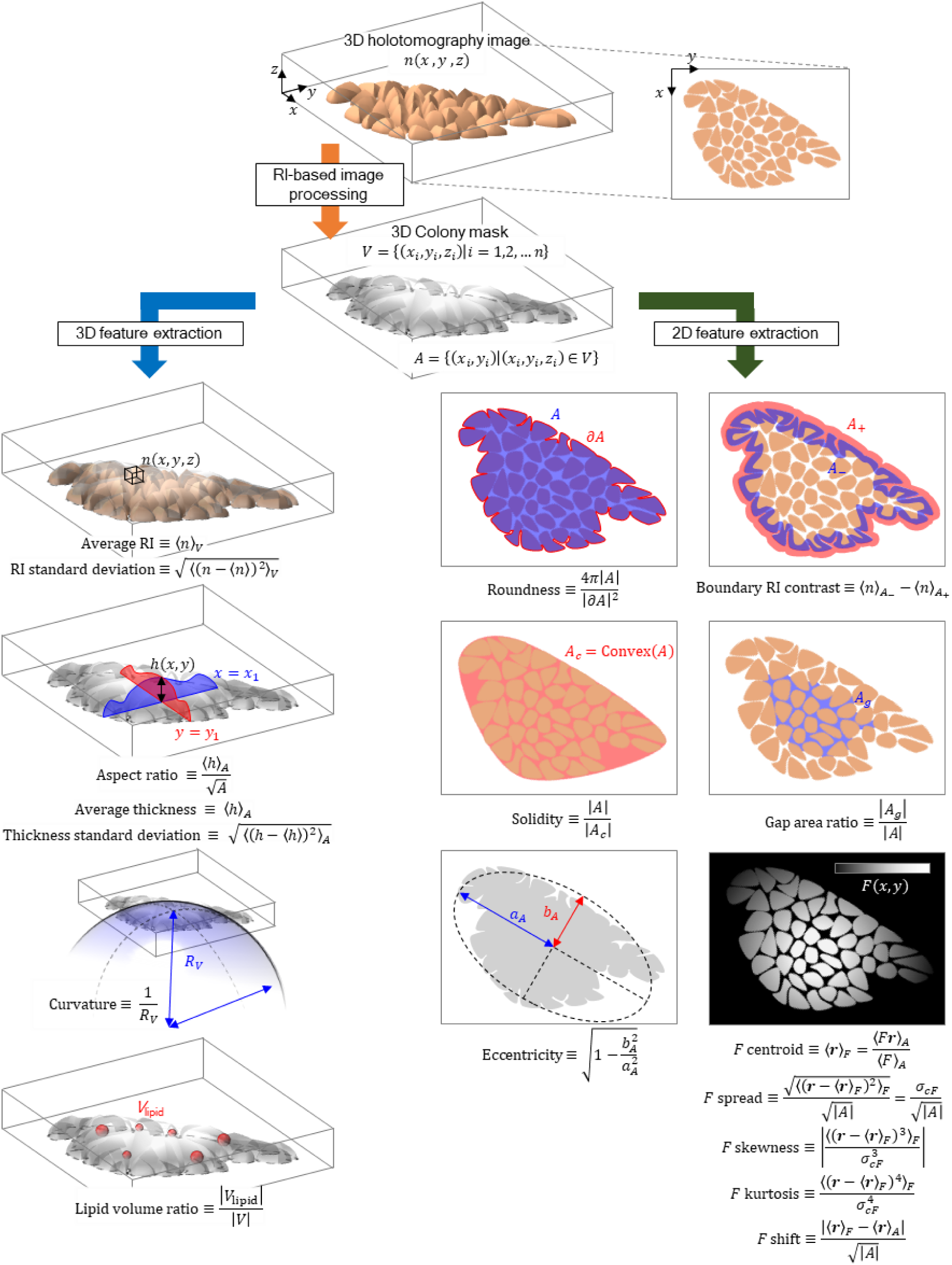
Graphical description of deriving quantitative hPSC colony properties from HT measurement. *N*: RI. *V*: 3D volume mask. *A*: 2D area mask. *H*: Thickness. *R_V_*: radius of the sphere fitting the upper surface of *V*. ∂*A*: 2D area boundary. *A_+_*: 2D area mask 2 μm outwards from ∂*A*. *A_-_*: 2D area mask 2 μm inwards from ∂*A*. *A_g_*: 2D area mask of intercellular gaps. Convex: convex hull construction. *A_A_*: Semi-major axis of the ellipse fitting *A*. *b_A_*: Semi-minor axis of the ellipse fitting *A*. *F* can be chosen among the spatially varying numerics including RI (*h*), thickness (*h*), dry mass, presence of lipid, and presence of gaps.

**Extended Data Fig. 9.**
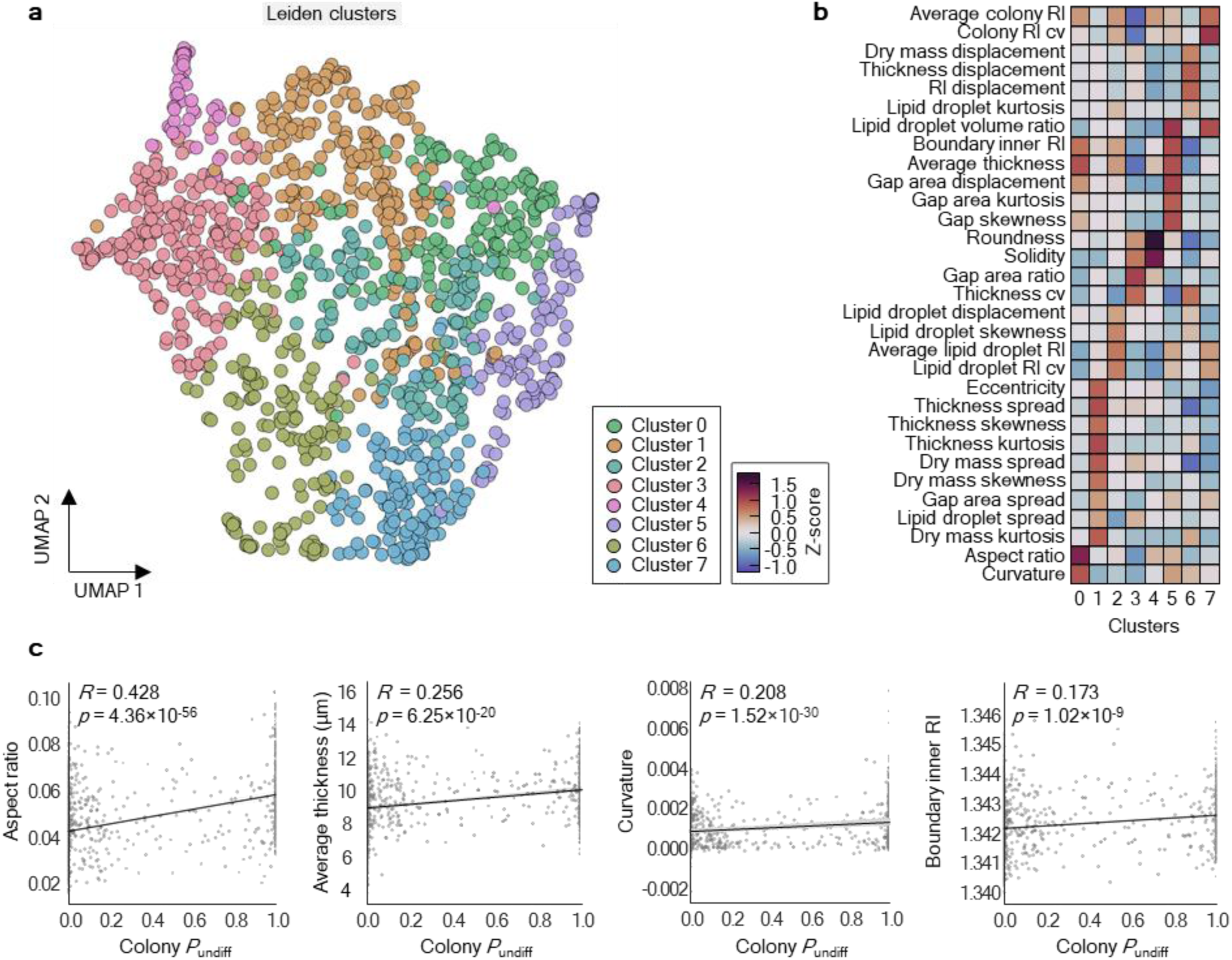
HT-derived hiPSC colony properties and their relation to the DeepHOPE result. **(a)** Distribution of HT-derived colony properties, visualised through UMAP embedding. **(b)** Leiden clustering (*k* = 8) by colony properties and heatmap of properties in clusters. **(c)** Linear fit between *P*_undiff_ and aspect ratio, average thickness, curvature and average RI MIP at the inner boundary of each colony. *R* and *p* refer to Pearson’s correlation coefficient and *p*-value respectively.

**Extended Data Fig. 10.**
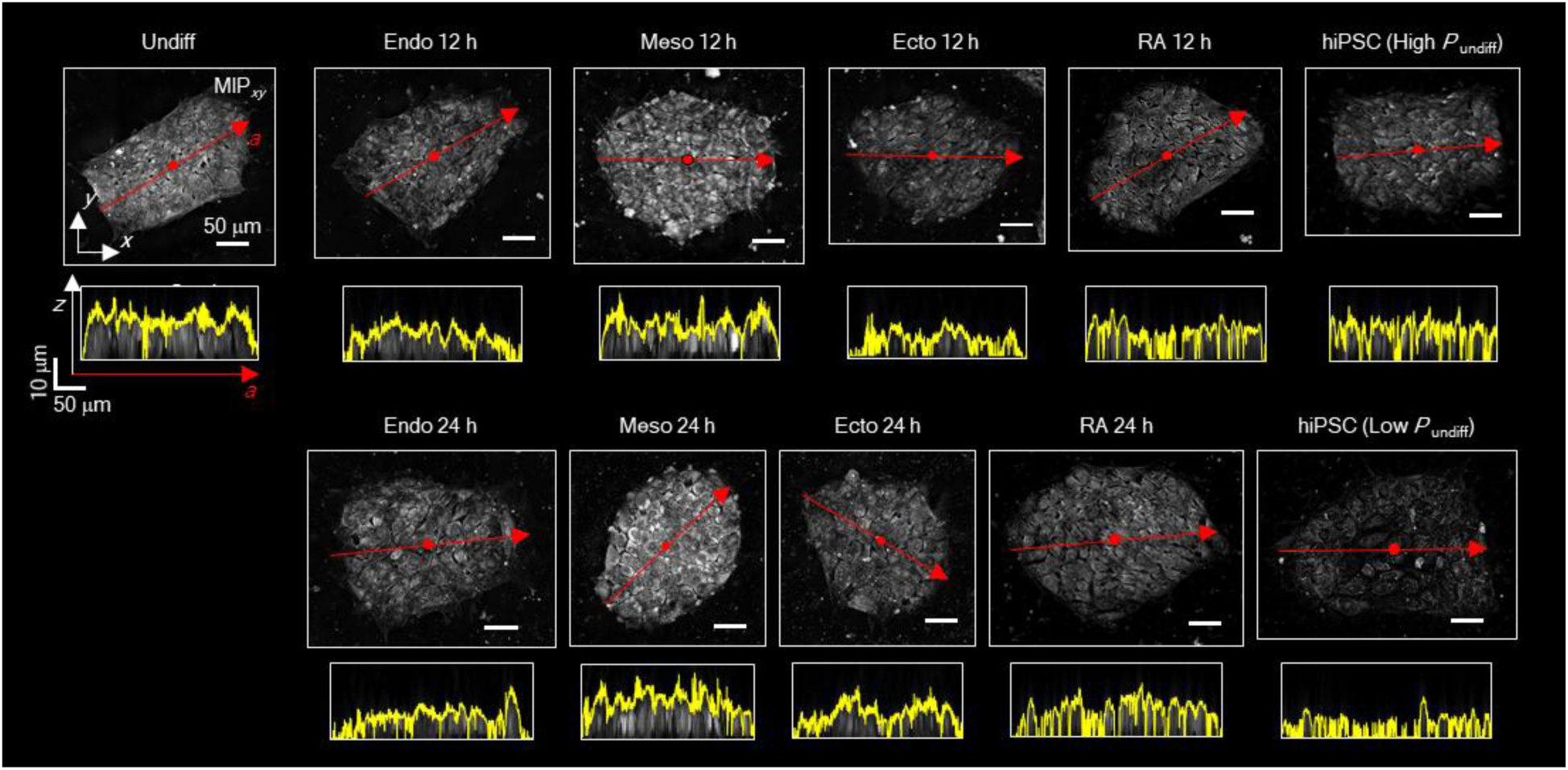
Topological diversity in hiPSC colonies under different differentiation status. Representative *xy* MIP and sectional view along the major axis (red arrow) of hiPSC colonies. The yellow line in each sectional view highlights the upper surface profile. Scale bar = 50 μm.

**Extended Data Fig. 11.**
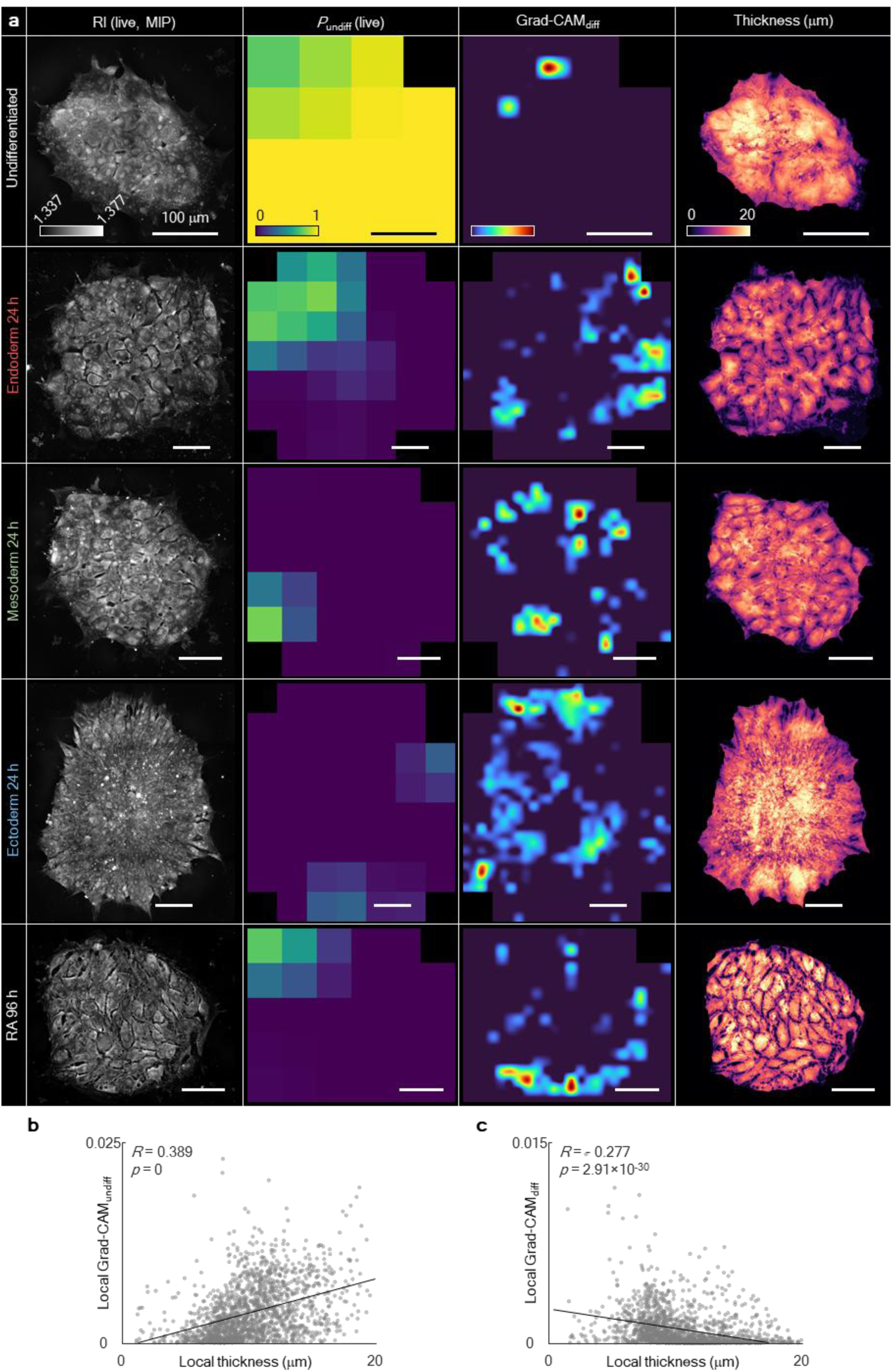
Visualisation of neural activation during DNN image processing of DeepHOPE. **(a)** Each column represents the HT measurement, probability map generated by DeepHOPE evaluation, Grad-CAM map for predicting *undifferentiated*, Grad-CAM map for predicting *differentiating* and thickness map. Scale bar = 100 μm. **(b** and **c)** Linear fit between the local colony thickness and local Grad-CAM score (B: class “*undifferentiated*”, C: class “*differentiating*”) across the GM trilineage dataset. *R* and *p* refer to Pearson’s correlation coefficient and *p*-value respectively.

